# Structure and dynamics of the contractile vacuole complex in *Tetrahymena thermophila*

**DOI:** 10.1101/2023.09.13.557576

**Authors:** Chao-Yin Cheng, Daniel P. Romero, Martin Zoltner, Meng-Chao Yao, Aaron P. Turkewitz

## Abstract

The contractile vacuole complex (CVC) is a dynamic and morphologically complex membrane organelle, comprised of a large vesicle (bladder) linked with a tubular reticulum (spongiome). CVCs provide key osmoregulatory roles across diverse eukaryotic lineages, but probing the mechanisms underlying the structure and function is hampered by the limited tools available for *in vivo* analysis. In the experimentally tractable ciliate *Tetrahymena thermophila*, we describe four proteins that, as endogenously tagged constructs, localize specifically to distinct CVC zones. The DOPEY homolog Dop1p and the CORVET subunit Vps8Dp localize both to the bladder and spongiome but with different local distributions that are sensitive to osmotic perturbation, while the lipid scramblase Scr7p co-localizes with Vps8Dp. The H^+^- ATPase subunit Vma4 is spongiome-specific. The live imaging permitted by these probes revealed dynamics at multiple scales including rapid exchange of CVC-localized and soluble protein pools vs. lateral diffusion in the spongiome, spongiome extension and branching, and CVC formation during mitosis. While the association with *DOP1* and *VPS8D* implicate the CVC in endosomal trafficking, both the bladder and spongiome are isolated from bulk endocytic input.

**Summary statement:** In the ciliate *Tetrahymena thermophila*, four proteins are shown to provide markers for different zones of the contractile vacuole complex. They shed light on its formation and maintenance by enabling *in vivo* analysis of its dynamics.

## Introduction

Though essential for life, water also endangers any cell that cannot offset diffusional water uptake across its plasma membrane. Inward water diffusion occurs whenever the tonicity of the cell cytoplasm exceeds that of the extracellular fluid, and this can result in catastrophic swelling for cells that lack a rigid cell wall (Ritter et al., 2021). This challenge existed during the evolution of many unicellular protists adapting to fresh-water environments, and is today reflected in the phylogenetically widespread presence of osmoregulatory organelles called contractile vacuole (CV) complexes (CVCs) (Allen, 2000, Jimenez et al., 2022, Patterson, 1980). The CVC consists of one or more large vesicles, sometimes called cisternae or bladders (a name we will use) which are connected to membrane tubules that extend as a network (spongiome) into the cytoplasm (Allen and Naitoh, 2002). In current models, water absorbed into the spongiome from the cytoplasm fills the expanding CV, which subsequently contracts during the phase when its contents are expelled through a transient opening in the plasma membrane (Plattner, 2015). As befits an osmoregulatory organelle, the periodicity of the contractile cycle is sensitive to the extracellular tonicity (Allen, 2000, Docampo et al., 2013, Heuser et al., 1993, Gabriel et al., 1999).

CVCs were first recognized as prominent cellular features more than two centuries ago but our current understanding of mechanisms underlying cyclic expansion/contraction is still fragmentary, including of the mechanics of contraction itself (Tani et al., 2001, Tani et al., 2000, Tani et al., 2002, Naitoh et al., 1997, Spallanzani, 1776). The most detailed molecular studies have been pursued in *Dictyostelium discoideum*, *Paramecium* (species *multimicronucleatum* and *tetraurelia*), and the Trypanosomid *T. cruzi*, which represent three highly unrelated lineages (Amoebozoa, Alveolata, and Rhizaria, respectively). In all three, water transfer from the cytosol into the CVC lumen involves the activity of proton-pumping vacuolar ATPases in the organellar membrane (Fok et al., 1995, Gronlien et al., 2002, Temesvari et al., 1996). However, because the CVC lumen is not detectibly acidified (Stock et al., 2002a), it is believed that the proton gradients produced by these pumps are harnessed to produce coupled gradients of other ions, which are more directly involved in transport of water (Jimenez and Docampo, 2015) and other small molecules (Stock et al., 2001, Stock et al., 2002a, Stock et al., 2002b, Heuser et al., 1993). Bladders may also undergo fusion with cytoplasmic organelles called acidocalcisomes that independently accumulate water-drawing ions (Rohloff et al., 2004, Marchesini et al., 2002, Montalvetti et al., 2004). However, the details are unclear and may differ between organisms, between which there are also notable morphological differences (Allen and Naitoh, 2002). For example, in Trypanosomes, the bladder is localized by attachment via an electron-dense structure to the flagellar pocket, in which transient pores are posited to form during bladder emptying (Girard-Dias et al., 2012). A similar arrangement exists in Paramecium, except that the bladder localization is via linkage to stable plasma membrane pores via dedicated cytoskeletal connections (Allen and Naitoh, 2002). In contrast, in Dictyostelium both the number and the position of bladders is variable, and their emptying appears to involve kiss-and-run fusion at contingent sites on the plasma membrane (Becker et al., 1999, Essid et al., 2012, Heuser, 2006).

Similarly, the anatomy of the tubular network surrounding the bladder, the spongiome, as well as the connections between the bladder and spongiome, display significant differences between these organisms (Allen and Naitoh, 2002). In Dictyostelium, both the bladder and spongiome are rich in vacuolar ATPase, and the two structures may interconvert during contractile cycles (Gerisch et al., 2002). In *Trypanosoma cruzi* the V-ATPase appears concentrated in the bladder (Ulrich 2011 PLOS One), while the Paramecium V-ATPase is strictly concentrated in a sub-domain of the spongiome, called “decorated” (Fok et al., 1995). The Paramecium CVC, which altogether appears to be more highly structured than in Dictyostelium or Trypanosoma, also includes a third distinct zone comprised of undecorated tubular arms, that links the bladder and decorated spongiome (Ishida et al., 1996, McKanna, 1976). The arms appear to become cyclically isolated from the bladder in a way that limits fluid backflow during contraction (Tominaga et al., 1998b, Gronlien et al., 2002, Tani et al., 2002). A similar dynamic arrangement has been described in Trypanosomids (Linder and Staehelin, 1979).

Contractile vacuole-associated proteins have been identified in Dictyostelium, Trypanosomes, and Paramecium, but only a small subset of these appeared uniquely localized to the CVC (Plattner, 2013, Bush et al., 1994, Du et al., 2008, Manna et al., 2023, Plattner, 2015, Schonemann et al., 2013, Ulrich et al., 2011, Gerald et al., 2002, Gabriel et al., 1999). This subset has provided key tools both for live imaging of CVC dynamics in the form of fluorescently-tagged transgenes, as well as for advancing informative candidates for functional studies (Becker et al., 1999, Du et al., 2008, Gabriel et al., 1999, Gerisch et al., 2002, Harris et al., 2001). In a prior study, we identified such a candidate in the ciliate *Tetrahymena thermophila*, as part of a survey of CORVET complexes in this organism (Sparvoli et al., 2020). Tetrahymena, like Paramecium, belongs to the Oligohymenophorean subgroup of ciliates, but the two genera diverged from one another hundreds of millions of years ago (Warren et al., 2017). VPS8D encodes one subunit of a hetero-hexameric complex whose homologs in other organisms are involved in vesicle fusion in the endolysosomal pathway (van der Beek et al., 2019b). In Tetrahymena, one of six such complexes appeared to localize to the CVC (Sparvoli et al., 2020). Though the CVC in Tetrahymena species was historically the subject of both physiological and ultrastructural studies (Elliott and Bak, 1964), Vps8Dp (the protein product of the *VPS8D* gene), together with a small set of Rab GTPases (Bright et al., 2010), were the first potential protein markers specific to this compartment. Recently, other potential candidates were advanced in genetic studies that uncovered two genes, *DOP1* and *VMA4*, linked with CVC-related phenotypes (Cheng et al., 2016). *DOP1* belonged to the conserved DOPEY family, of which members in yeast and animals were associated with membrane trafficking (Moliere et al., 2022b). *VMA4* encoded the E-subunit of a V-ATPase complex (Collins and Forgac, 2020).

In this manuscript, we provide evidence that Vps8Dp, Dop1p, and Vma4p, together with a novel predicted lipid scramblase Scr7p, are bona fide CVC proteins in *T. thermophila*. Vps8D and Dop1p also have large dispersed cytoplasmic pools, which undergo exchange with the CVC-localized cohort. By exploiting fluorescently-tagged copies of these proteins for live cell imaging, we developed models for both the anatomy and the contractile cycle of the CVC in this organism. Cells expressing the tagged proteins also provided new insights into the response of the CVC to osmotic stress, and CVC duplication during cell division. Notably, although all four proteins localize to the CVC, they occupy different zones. Dop1p and Vps8Dp both localize to the bladder periphery, but Vps8D is shifted distally. This difference between localization of Vps8Dp and Dop1p becomes more pronounced under hypo-osmotic stress, supporting the idea that the proteins occupy two functionally distinct regions. In addition, Vps8Dp and Dop1p localize to the spongiome, but here too their distribution is not identical. Instead, the two appear locally concentrated within distinct but overlapping zones. Scr7p localization coincides with that of Vps8Dp, while Vma4p is strictly localized to the spongiome but not the bladder.

## Results

### 1. Dop1p, a protein in the DOPEY family, localizes to the CVC bladder and spongiome

*T. thermophila* contains a prominent CVC, whose structure based on past studies consists of a central bladder that is bridged with the plasma membrane via pores, generally two in number. The bladder is surrounded by a tubulovesicular spongiome. This structure is shown in Fig. 1A. By standard light microscopy only the bladder is visible, as a round structure in the cell posterior. The bladder goes through cycles of collapse and expansion on a time scale of tens of seconds (Fig. 1B, top row). As previously reported, cells lacking the *DOP1* gene instead develop an enormously enlarged bladder which grows at least in part by undergoing fusion with other large vesicular structures, and only undergoes cyclic collapse on a time scale of minutes (Fig. 1B, bottom row (Cheng et al., 2016)). To ask whether the protein product of *DOP1* is directly associated with the contractile vacuole, we targeted the Macronuclear *DOP1* locus to endogenously tag the Dop1p C-terminus with mNeon. (The micronuclear germline *DOP1* locus, which was not modified by our approach, is silent in vegetative cells (Cassidy-Hanley et al., 1997). Cells expressing Dop1p-mNeon showed bright fluorescence in the cell posterior. In cross sectional views, the bright fluorescent signal was tightly localized to the periphery of the CV bladder, from which it extended both anteriorly and posteriorly as a stripe tracing the cell cortex (Fig. 1C). In cells that are rotated 90° around their long axis, relative to the perspective in Fig 1C, Dop1p signal is again most concentrated at the CV bladder and extends in an irregular reticulum that radiates in all directions (Fig. 1D). In near-tangential views the fluorescent signal appeared as two (or sometimes three) rings whose diameters were smaller than that of the bladder, and whose positions suggested their potential association with the pores bridging to the plasma membrane through which the bladder empties (Fig. 1E). To investigate this further, we took advantage of the fact that bundles of microtubules are known to emanate from the pore junctional zone linking the bladder and plasma membranes (Gaertig et al., 1995, Frankel, 2000), and thus the structures can be stained using anti-α-tubulin antibodies. In cells where we visualized both Dop1p and α- tubulin, the rings formed by the two proteins were concentric but not significantly overlapping (Fig. 1F and 1G). These images suggest that the junctional pore membrane may have a different composition from the bladder, and that Dop1p localizes preferentially to the latter. In addition to this concentrated fluorescence at the bladder and surrounding spongiome, Dop1p puncta were also lightly dispersed throughout the cell cytoplasm.

**Figure 1:**
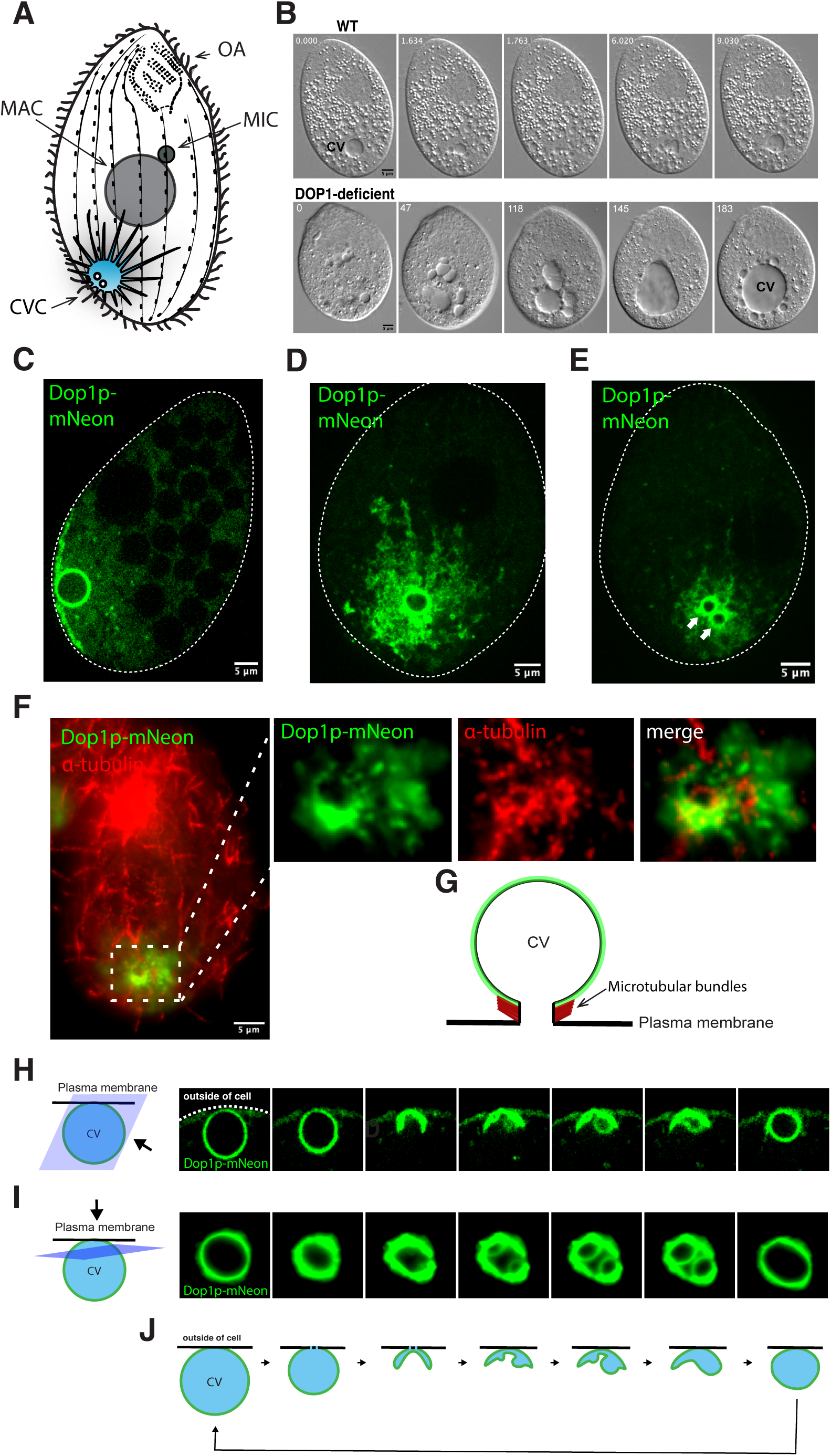
Dop1p is a live cell marker for the Tetrahymena CVC. A: CVC anatomy in Tetrahymena. A cartoon showing previously established features of the Tetrahymena CVC including one central bladder near the cell posterior, the surrounding tubulo-vesicular spongiome network, and two pores connecting the bladder to the plasma membrane. Also shown are the oral apparatus (OA) in the anterior end of cell, and both the micronucleus (MIC) and macronucleus (MAC) near the cell middle. The vertical rows represent the cytoskeletal tracks called primary meridians, along which the ciliary basal bodies are spaced. For clarity, the cilia themselves are only shown at the cell edges. B: As previously reported, cells deficient in *DOP1* show pronounced changes in the structure and dynamics of the bladder. Shown are five sequential DIC (differential interference contrast) images (individual frame for imaging was shown at t= 0s, 1.634s, 1.763s, 6.02s and 9.03s of WT images and for *DOP1*-deficient cells at t= 0s, 47s, 118s, 145s and 183s, respectively labeled in left-upper comers in the images from video footage of WT and *DOP1*-deficient cells (Cheng et al., 2016), demonstrating profound differences in the appearance and cycling time of the CV bladder in the mutant cells. Whereas the WT cell bladder shown goes through a contractile cycle in ∼10 seconds, the mutant cell bladder does not undergo a contraction over the 3 minute interval shown but instead continued to expand. C-E: Expression of Dop1p endogenously tagged with mNeon. Cells from growing cultures were immobilized for imaging using CyGEL, as described in Materials and Methods. Videos were captured using a Marianas spinning disc confocal microscope with the fastest speed model. C. The tagged protein localizes primarily at the bladder and along structures that extend from the bladder beneath the plasma membrane. Shown is one frame from a time-lapse video with 0.1 sec frame intervals, focused near the cell mid-section (Movie 1). D: Imaging of a cell, rotated along its long axis compared to that in C, so that the CVC is near the top surface of the cell. In this orientation, the structures underlying the plasma membrane are seen to constitute a reticular network. Shown is one frame from a time-lapse video with 0.51 sec frame intervals (Movie 2). E: Imaging of a cell oriented as in D, but with the focal plane very near the plasma membrane. In this tangential focal plane, Dop1p localizes to the circular area which may be related to the two pores. Shown is one frame from a time-lapse video with 0.51 sec frame intervals (Movie 3). F: Imaging of a fixed cell expressing Dop1p-mNeon with immunofluorescent staining of α-tubulin, in a focal plane very near the cell surface. The area within the dotted white line, containing the two CV pores, is shown enlarged in the right panels with both the individual and the merged images. Dop1p does not superimpose on the microtubular bundles of the pores. Images were taken with a Zeiss Axio Observer 7 system. G: Cartoon showing the known position of microtubules that emanate from the sides of the CV pore. H: Live imaging of the CVC in a Dop1p-mNeon expressing cell. The focal plane is roughly at the midpoint of the expanded bladder, viewed from the angle shown by the arrow. The seven images were extracted from a video capturing a contractile cycle. (Movie 4). Cells from growing cultures were immobilized for imaging using CyGEL, as described in Materials and Methods. Videos were captured using a Marianas spinning disc confocal microscope with the fastest speed model. I: Live imaging of the CVC in a Dop1p-mNeon expressing cell. Here the focal plane is tangential to the expanded bladder, close to the edge where it contacts the plasma membrane, as viewed from the angle shown by the arrow. The seven successive images were extracted from a video capturing a contractile cycle (Movie 5). Cells were immobilized by applying slight pressure to the cover slip (described in Materials and Methods) and imaged using a Zeiss Axio Observer 7 system with the fastest speed model to capture each frame among multiple exposures. J: A model for the contractile cycle of the bladder. The model to illustrate the polarized collapse and refilling of the bladder is based on live images as exemplified in panels H and I.

The availability of Dop1p as a bright and relatively compartment-specific marker for the CVC offered a new addition to the cell biology reagents available for Ciliates, and it allowed us to monitor aspects of the contractile cycle in live cells. In optical sections corresponding roughly to the midpoint of an expanded bladder, one sector of the circumference of the expanded bladder is apposed to the cell periphery. That sector appears to be relatively stable when the CV contracts, while the rest of the bladder apparently collapses down upon it. This results in a brightly fluorescent crescent, which in three dimensions would appear as a lens. That is, the replacement of a CV sphere by a lens can be most easily explained by directional collapse of the bladder so that the cortex-distal zone draws close to the cortex-proximal zone (Fig. 1H). This model suggests that the PM-adjacent zone of the bladder is selectively stabilized, which may involve the aforementioned microtubules that emanate from the CV-plasma membrane junctions (Fig. 1G). The lens serves as a platform for the refilling of the bladder. As seen in cross section, refilling involves the emergence from the crescent of what appear as blebs, which subsequently expand (Fig. 1H).

The same cycle could also be viewed tangentially in living cells, in optical sections just below the surface that are roughly coplanar with the plasma membrane in the region of its junction with the CV (see cartoon in Fig. 1I). From this perspective, the ring corresponding to the bladder periphery fluctuates in fluorescence intensity over the contractile cycle but shows only minor fluctuations in size (Fig. 1I). These images are consistent with the idea that the crescent seen in the 3^rd^ and 4^th^ panels of Figure 1H is a cross-section of a lens-shaped compartment that results from collapse of the originally spherical bladder (Fig. 1I). From the same tangential perspective, the blebbing seen in the orthogonal view (Fig. 1H, 4^th^ panel) manifests as twin circles (Fig. 1I, 4^th^ panel), suggesting that this phase of membrane expansion may be structured by the pores. A model for the contractile cycle of the bladder based on these complementary perspectives is shown in Figure 1J.

### 2. Using Dop1p as a live marker allows visualization of contractile vacuole duplication

During cell division, a subset of structures present in a cell must be duplicated and/or partitioned so that they are inherited by both daughters. In *T. thermophila* cell division (Fig. 2A), key features of the cell are duplicated along an anterior-posterior axis prior to cytokinesis (Cole and Gaertig, 2022). These include the CV pores, in which a new anterior pair appears at a stage when the Micronucleus but not Macronucleus has undergone mitosis. The pre-existing CV remains in the posterior while a second CV forms in the anterior and at the same relative position on the cell cortex (Ng, 1977, Ng and Frankel, 1977, Frankel, 1992, Frankel, 2000).

**Figure 2:**
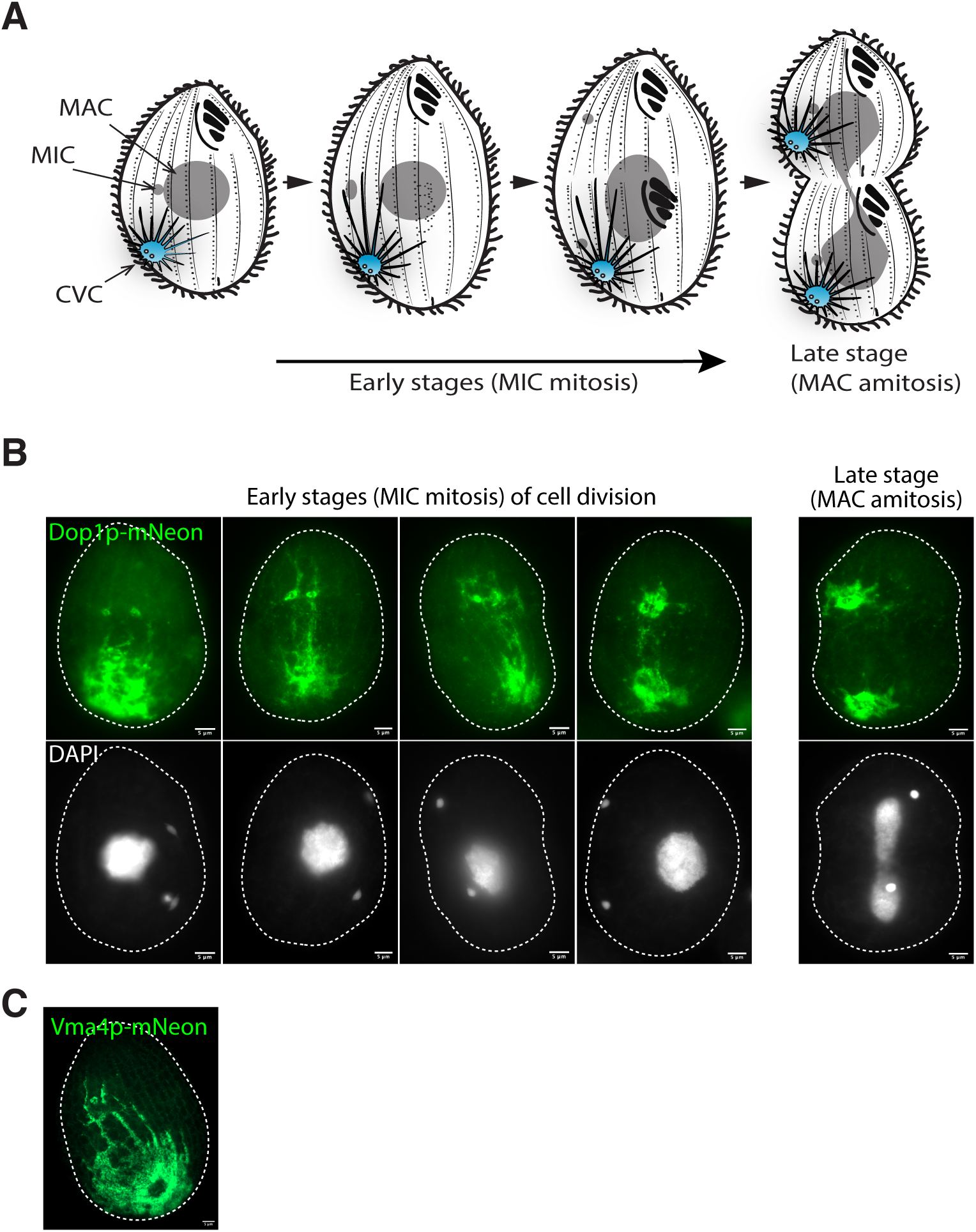
Dop1p-based imaging of CVC duplication. A: Cell division in *Tetrahymena thermophila*. Micronuclear (MIC) mitotic division is followed by macronuclear (MAC) elongation and amitotic division. Unique cellular features, including the CVC and oral apparatus, are generated as second copies in the anterior half of the elongating cell, which will become the daughter upon cytokinesis. B: Imaging of the CVC in Dop1p-mNeon-expressing dividing cells, that were fixed and stained with DAPI. The first four images are of cells at early stages in cell division, as judged based on nuclear morphology, while the 5^th^ image captures a later stage. The newly forming CVC in the anterior is first visible as two Dop1p-labeled circular structures that may represent incipient bladders. Dop1p also appears to be selectively concentrated along a subset of cytoskeletal meridians that extend from the parental to the new CVC. Growing cells were fixed and stained as described in Materials and Methods. Images were taken with a Zeiss Axio Observer 7. C: Imaging of the CVC in a fixed Vma4p-mNeon-expressing dividing cell. The newly forming CVC in the anterior is recognizable by the initiation of a Vma4p-labeled spongiome that appears connected along a subset of cytoskeletal meridians with the parental CVC. Growing cells were fixed as described in Materials and Methods. Images were taken with a Zeiss Axio Observer 7.

We followed this process in cells expressing Dop1p-mNeon. In particular, while previous workers had focused exclusively on the pores themselves as prominent cortical landmarks in fixed cells, the availability of a CV marker allowed us to visualize other compartments and in live cells. At an early stage in the process when only the Mic has undergone mitosis, Dop1p signal extends anteriorly from the maternal CV beyond the elongating cell midline, to two circular structures which are located at the cortical position that will be occupied by the newly forming CV (Fig. 2B). These circular structures may be related to the pores but their diameter (2-3µM) is too large to correspond to mature pores. We favor the idea that the Dop1p-labeled structures instead reflect the beginning of bladder formation, an idea more consistent with the bladder localization data in Fig. 1G.

Many of the Dop1p-labeled puncta and Vma4p-labeled reticulum seen in dividing cells are aligned along the cytoskeletal “ribs”, called meridians, that underlie the Tetrahymena cortex (Fig 2B, 2C and S4) (Frankel, 2000). Interestingly, *DOPEY* family members (called DOPEY) in other organisms can recruit kinesin, a cytoskeletal-based motor protein (Mahajan et al., 2019). We therefore wondered whether *T. thermophila* Dop1p might associate with the cytoskeleton by a similar mechanism. A pilot immuno-isolation of epitope-tagged *DOP1*, expressed at the endogenous locus, was analyzed by mass spectrometry to identify potential interacting proteins. We detected two predicted scavenger mRNA decapping proteins, a TATA-binding protein interacting protein, and a predicted polytopic transmembrane protein (Fig. S2, Table S4 and S5). Based on the Conserved Domain Search (NCBI), the polytopic protein may contain a PHM7_cyt Super family domain, which is a characteristic of ion transporters including osmosensitive or mechanosensitive ion channels (Wang et al., 2023). To ask whether this protein associates with Dop1p at the CV bladder, we endogenously tagged it with mNeon. At steady state, the tagged protein was not visible at the CVC but instead appeared localized to peripheral structures that are likely to be the Golgi, as well as heterogeneous mobile compartments deeper in the cytoplasm (Fig. S3 and Movie 17). Given this lack of localization to the CV, together with the lack of an obvious connection to RNA metabolism, we did not further pursue these pulldown-defined candidates.

### 3. Vps8Dp, a CORVET complex subunit, shows punctate localization to the CVC

CORVET is a widely conserved hetero-hexameric complex, characterized most extensively in yeast and animals, that facilitates vesicle-vesicle tethering and fusion in endosomal trafficking (Balderhaar and Ungermann, 2013). We previously established that *T. thermophila* expresses six distinct CORVET complexes, called A through F, and we found that the D complex was likely to associate with contractile vacuoles by localizing its Vps8 subunit (Vps8Dp) (Sparvoli et al., 2020). To ask how Vps8Dp localization compared with that of Dop1p, we analyzed cells expressing *VPS8D* endogenously tagged with mNeon. Consistent with previous results, Vps8Dp-mNeon appeared to localize both around the CV bladder and to the spongiome (Fig. 3A and 3B), and like Dop1p also revealed the twin connections between the bladder and plasma membrane pores (Fig. 3B, third panel). However, whereas in live cell imaging the Dop1p fluorescent signal over the bladder and reticulum surfaces appeared relatively continuous, Vps8Dp labeling was distinctly punctate on all structures (Fig. 3B, 3C and 3D). In addition, heterogeneous puncta of Vps8Dp-mNeon were conspicuous throughout the cytoplasm.

**Figure 3:**
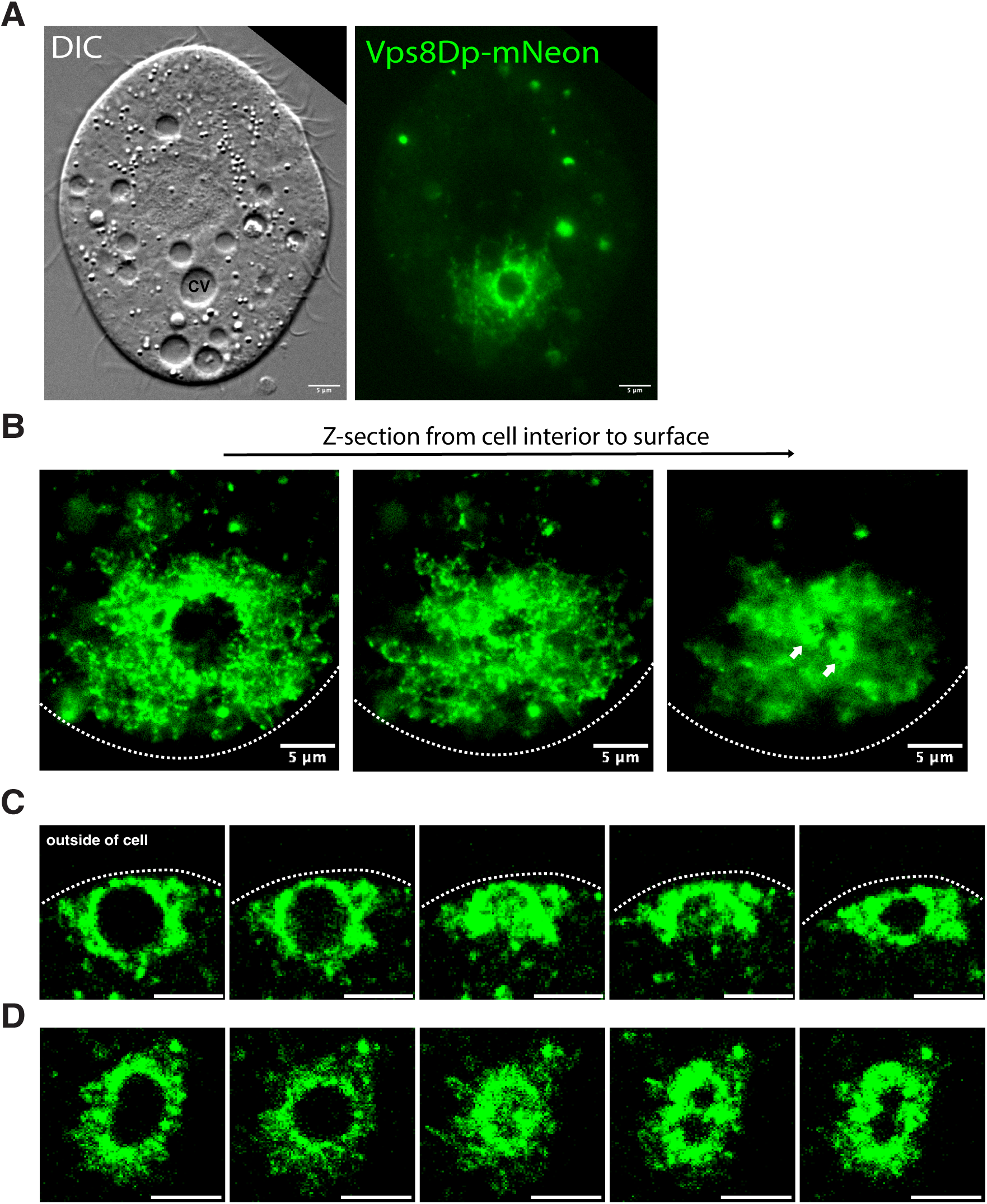
Vps8Dp, a subunit of CORVET, shows punctate localization throughout the CVC. A-D: Live imaging of the CVC in Vps8Dp-mNeon expressing cells. A. Left panel: Differential Interference Contrast (DIC) image of *Tetrahymena*; the CV bladder is labeled. Right panel: corresponding fluorescent image. The frame shown is from a time-lapse video with 2.73 sec frame interval (Movie 6A and 6B). B: Successive sections in a Z-stack. The first section corresponds roughly to the midpoint of the expanded bladder, and second and third sections are progressively closer to the bladder periphery and cell surface, where two pores are outlined. The time-lapse video (0.15 sec frame interval) from which these were taken is Movie 7. Both A and B were captured using a Axio Observer 7 system. C: Cross-sectional view of contractile cycle (perspective equivalent to Fig 1H). The video from which these five successive images were extracted is Movie 8. D: Tangential view of contractile cycle (perspective equivalent to Fig 1I). The video from which these five successive images were extracted is Movie 9. For both C and D, cells from growing cultures were immobilized using CyGEL and imaged using a Marianas spinning disc confocal microscope with the fastest speed model to capture each frame among multiple exposures.

As with cells expressing tagged Dop1p, the contractile cycle could be visualized in Vps8Dp-mNeon cells. Analysis of these videos confirmed that contraction is asymmetrical, with the bladder collapsing toward the plasma membrane (Fig. 3C). The images are consistent with the crescent/lens intermediate discussed above, but this structure appears less well-defined due to the fact that while Dop1p tightly traces the contour of the bladder membrane, Vps8Dp puncta are irregularly distributed near the bladder membrane (Fig. 3B, middle panel). When the contractile cycle is viewed along the axis of the pores, and just beneath the plasma membrane, the bladder is deformed but without undergoing dramatic contraction, consistent with imaging in Dop1p-mNeon cells (Fig. 3C, 3D and Fig 1).

### 4. *VMA4* localizes to the CVC reticulum but not to the bladder

Though the CV bladder and spongiome both contain Dop1p and Vps8Dp, these two compartments are likely to serve non-identical functions and therefore differ in their molecular composition. Vacuolar ATPases have been localized to a spongiome in both the amoebozoan *Dictyostelium discoideum* and the ciliate *Paramecium tetraurelia* (Fok et al., 1995, Heuser et al., 1993, Fok et al., 1993, Nolta et al., 1993). In *T. thermophila*, the vATPase E-subunit-encoding gene, *VMA4*, is a potential CVC determinant (Cheng et al., 2016). To test this, we disrupted all Macronuclear copies of *VMA4*. The resulting Δ*vma4* cells showed greatly slowed cycles of CV contraction (Fig. 4A). To ask whether Vma4p localizes to the CVC, we integrated the mNeon tag at the endogenous *VMA4* locus. The fluorescent signal was visible only at the CVC and exclusively within the spongiome. That is, unlike Vps8Dp or Dop1p, Vma4p was not concentrated at the bladder periphery (Fig. 4B and 4C). Imaging of the edges of the spongiome in live cells showed that Vma4p-labeled tubules show dynamic extension, retraction, and branching (Fig. 4D). The localization of Vma4p to the spongiome but not the bladder underscores the functional difference between those compartments.

**Figure 4:**
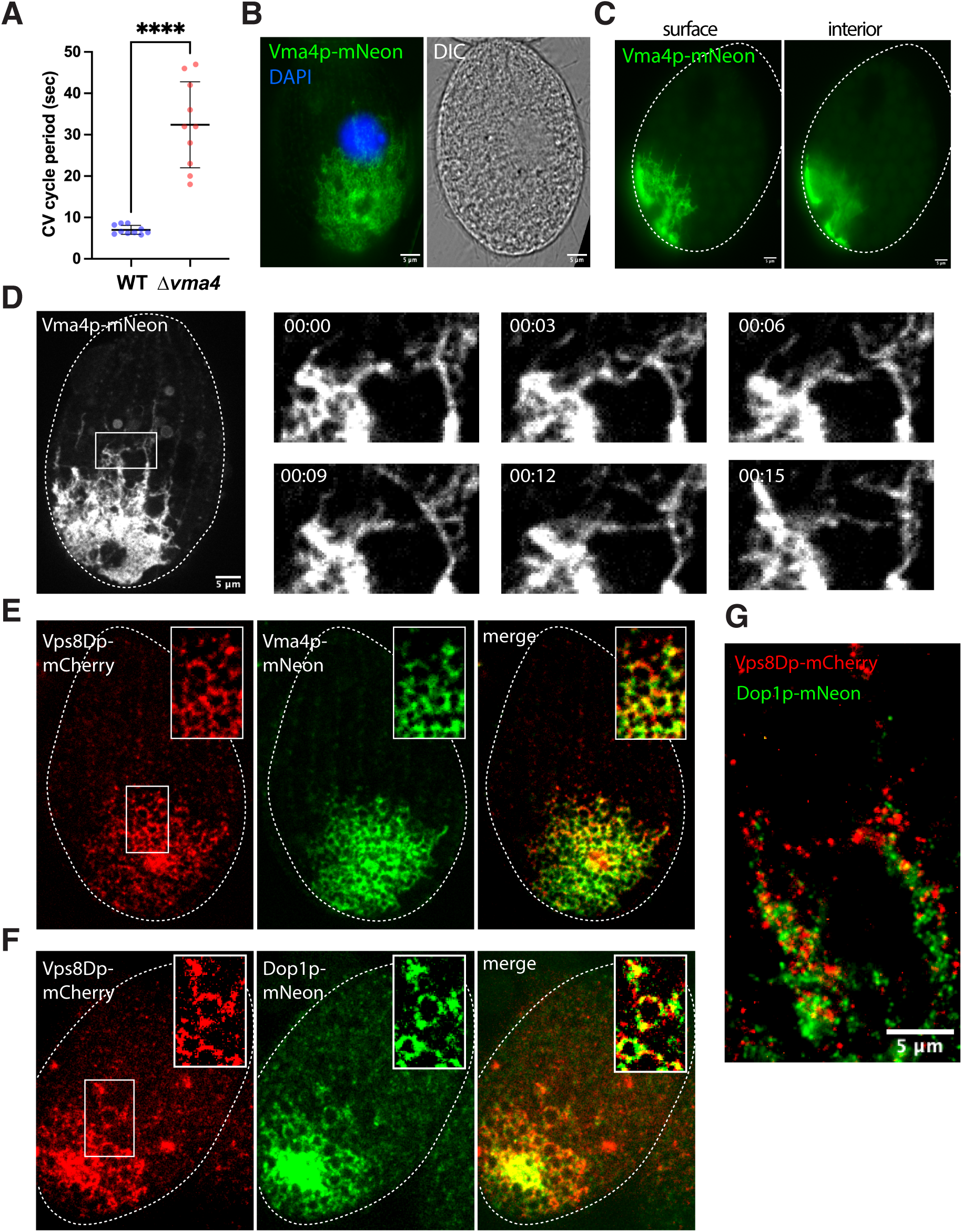
Vma4p is a CVC protein that localizes exclusively to the spongiome. A: The period of the contractile cycle was measured for cells from growing cultures of WT or Δ*vma4* (complete knockout of *VMA4*) (n=10 for each culture). The mutant cells showed a roughly 3-fold lengthening of the contractile cycle. The data were plotted using GraphPad Software Prism and using a two-tailed *t*-test. Individual data points and the mean±s.d. is shown. The difference between WT and Δ*vma4* is significant, *P*-value <0.0001. B: Vma4p localizes to the spongiome, but not the bladder. Growing cell expressing Vma4p-mNeon were fixed, and the nuclei stained with DAPI. A single cell is shown in the fluorescence (left) and DIC (right) channels. The Vma4p-labeled spongiome extends 10-20µM from the bladder periphery. Images were taken with a Zeiss Axio Observer 7 system. C: Vma4p-labeled tubules are concentrated near the plasma membrane, as shown in this side view of the CVC. The two panels are extracted from a Z-stack captured during live imaging (Movie 10), and show surface (left) and cell interior (right) sections. Images were taken with a Zeiss Axio Observer 7 system. D: Vma4p-labeled tubules show dynamic extending and branching. Left panel: Live imaging of Vma4p-labeled tubules. Right panels: magnified view of the boxed area in the left panel, showing six consecutive images (each individual image was selected with the 3 second interval, respectively labeled in left-upper comers in each image, extracted from a video (Movie 11). Images were taken using a Marianas spinning disc confocal microscope. E: Analysis of cells expressing Vps8Dp-mCherry and Vma4p-mNeon. Vps8Dp (left panel) and Vma4p (center panel) show extensive overlap over the entire spongiome (right panel, merge). The two proteins may have different local distributions along tubules (magnified insert in the three panels). Growing cells co-expressing Vps8Dp-mCherry and Vma4p-mNeon were fixed at low temperature to optimize spongiome preservation (as described in Materials and Methods). F: Analysis of cells expressing Vps8Dp-mCherry and Dop1p-mNeon. Vps8Dp (left panel) and Dop1p (center panel) show extensive overlap over the bladder and spongiome (right panel, merge). The two proteins may have different local distributions (magnified insert in the three panels). Growing cells co-expressing Vps8Dp-mCherry and Dop1p-mNeon were fixed at low temperature to optimize spongiome preservation (as described in Materials and Methods). Images of E and F were taken with a Marianas spinning disc confocal microscope. G: Expansion microscopy analysis of cells expressing Vps8Dp-mCherry and Dop1p-mNeon. A merged image shows the two proteins have different local distribution along the tubules. Growing cells co-expressing Vps8Dp-mCherry and Dop1p-mNeon were fixed at low temperature to optimize reticulum preservation and processed to expand the cell ∼10-fold, as described in Materials and Methods. Image was taken with a Zeiss Axio Observer 7 system.

The Vma4p-mNeon signal appeared evenly distributed along spongiome tubules, different from either Vps8Dp or Dop1p. To ask whether the same tubules contain Vma4p, Dop1p, and Vps8Dp, we expressed Vps8Dp-mCherry in pairwise combination with either Vma4p-mNeon or Dop1p-mNeon (Fig. 4E and 4F). In both cases, the images are consistent with the idea that all proteins are present and show overlapping distributions throughout the spongiome, but that there are local differences in their intensities (Fig. 4E and 4F), potentially reflecting membrane domains. These conclusions were consistent with expansion microscopy images of cells co-expressing Vps8Dp-mCherry and Dop1p-mNeon (Fig. 4G).

### 5. Proteins at the bladder periphery occupy subtly different zones

Since distinct CVC proteins appear differentially distributed within the spongiome, we wondered whether the same might be true of the bladder membrane. To investigate this possibility, we asked whether Dop1p and Vps8Dp differed in their distributions at the bladder periphery. We co-expressed Dop1p-mNeon and Vps8Dp-mCherry from their endogenous loci, and then traced the intensity of each signal along a vector drawn outward from the center of the bladder (Fig. 5A). This analysis revealed that the peak of Dop1p was proximal to that of Vps8Dp (Fig. 5B and 5C). This could be explained if Dop1p is bound at the bladder membrane while Vps8Dp is associated with a peripheral compartment, such as bladder-tethered vesicles. The Vps8Dp compartment appears less spatially restricted than the Dop1p compartment, since the intensity profile of Dop1p showed a narrow peak relative to that of Vps8Dp (Fig. 5B). We did the same pairwise comparison between Vps8Dp and Vma4p, the latter shown above to be associated with the spongiome, and as expected we found that Vma4p in such radial plots is yet further spaced from the center (Fig. 5D-F).

**Figure 5:**
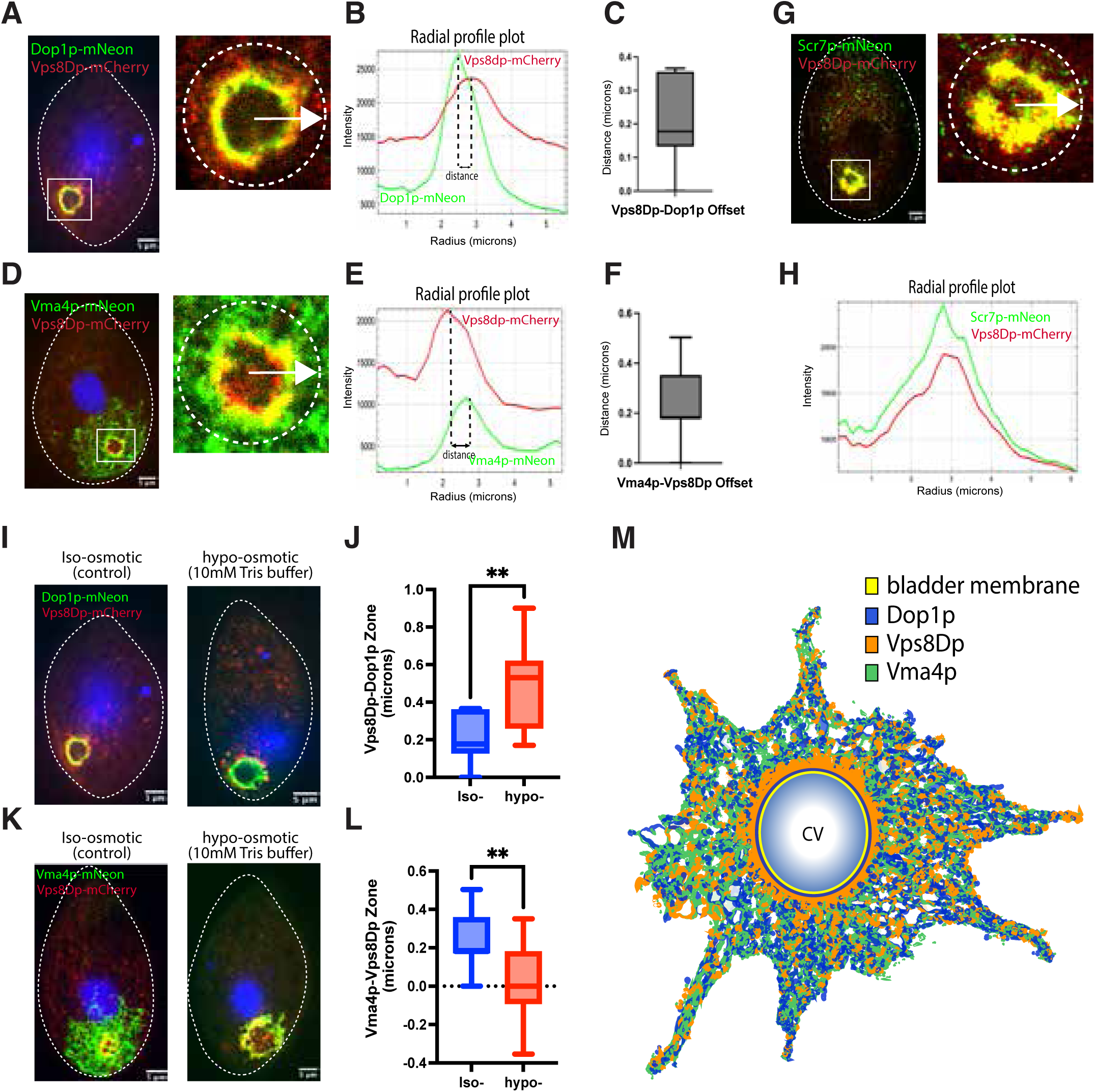
Finer mapping reveals distinct protein distributions at the bladder periphery and these distributions are sensitive to osmotic challenge. A-C: Vps8D is radially shifted with respect to Dop1p at the CV periphery. A: Left panel: growing cell co-expressing Dop1p-mNeon and Vps8Dp-mCherry. The boxed area, which includes the CV bladder, is enlarged in the right panel. The fluorescent intensity profiles of the two fluorophores along the radial arrow shown were determined using the FIJI image Radial Profile plugin. Images were taken with a Marianas spinning disc confocal microscope. B: The peak intensity of Vps8Dp is distal to that of Dop1p. The fluorescent intensity of Dop1-mNeon and Vps8Dp-mCherry was plotted along the vector in Panel A, drawn outward from the center of the CV bladder to the periphery. The distance between the peaks of Dop1p and Vps8Dp signals is shown. C: The zone distance between Vps8Dp and Dop1p. Fourteen additional images were analyzed as in Fig 5A,B. The combined data were plotted by GraphPad Software Prism and the mean±s.d. is shown. D-F: Vma4p is radially shifted with respect to Vps8D at the CV periphery. Left panel: growing cell co-expressing Vma4p-mNeon and Vps8Dp-mCherry. The analysis of this image was identical to that in Fig 5A,B. E: The peak intensity of Vma4p is distal to that of Vps8Dp. F: The zone distance between Vma4p and Vps8Dp. The experimental procedures and analytical methods used in Figure 5D-F are identical to those used in Figure 5A, 5B and 5C respectively. G-H: Scr7p co-localizes with Vps8Dp at the bladder periphery. G: Left panel: growing cell co-expressing Scr7p-mNeon and Vps8Dp-mCherry. Right panel: magnified image of boxed area containing the CV bladder. H: Scr7p and Vps8Dp show similar intensity profiles along a radial vector. The experimental procedures and analytical methods used in Figure 5G and 5H are identical to those used in Figure 5A and 5B respectively. I: Growing cells expressing Dop1p-mNeon and Vps8Dp-mCherry were kept in culture medium (left panel) or incubated for 10 min in 10mM Tris buffer (hypo-osmotic, right panel), as described in Materials and Methods. Fixed and DAPI-stained cells were imaged with a Marianas spinning disc confocal microscope. J: The zone distance between Dop1p-and Vps8dp increases under hypo-osmotic challenge. Fourteen images from control cells and thirteen images from the hypo-osmotic sample were used to measure the zone distance between Dop1p and Vps8Dp. The data were plotted by GraphPad Software Prism with statistical analysis via two-tailed *t*-test. The mean±s.d. is shown and *P*-values <0.05 indicate a significant increase in the spacing of the Vps8Dp and Dop1p peaks upon hypo-osmotic treatment. K: Cells expressing Vma4p-mNeon and Vps8Dp-mCherry. The experimental procedures and analytical methods are identical to those in Figure 5I. L: The zone distance between Vma4p and Vps8Dp decreases under hypo-osmotic challenge. Fourteen images from control cells and nine images from the hypo-osmotic sample were used to measure the zone distance between Vma4p and Vps8Dp. The mean±s.d. is shown and *P*-values < 0.05 indicate a significant decrease in the spacing of the Vma4p and Vps8Dp peaks upon hypo-osmotic treatment. M: A model for the distribution of Dop1p, Vps8Dp and Vma4p in the CVC. Dop1p associates tightly with the bladder membrane. Vps8Dp associates with a broader zone and is more distal. Vma4p primarily localizes in the spongiome and partially overlaps with Dop1p and Vps8Dp. All three proteins localize throughout the spongiome, but may have non-identical distributions.

Our results suggest that the bladder periphery may be conceived of as several concentric zones, for which Dop1p, Vps8Dp, and Vma4p are markers that can be used to map the distributions of additional proteins. For example, *SCR7* encodes a novel predicted lipid scramblase that localizes in part to the CVC (Fig. S5, Movie 19 and 20). When we co-expressed Scr7p-mNeon with Vps8D-mCherry, the two proteins showed near-complete co-localization (Fig. 5G, 5H and S5). Thus, Scr7p appears to localize to the same CVC zone as Vps8Dp. This finding also provides evidence that the distinct localizations seen for other pairs of proteins does not reflect a technical artefact, for example if the fluorescent protein tags induced homophilic clustering.

### 6. Distinct compartments of the CVC respond differentially to osmotic stress

Many observations of ciliates by simple light microscopy have revealed that osmotic stressors result in morphological and dynamic changes to the CV bladder (Allen and Naitoh, 2002). The proteins we have identified in *Tetrahymena* allowed us to extend these observations in live cells by visualizing the spongiome, as well as distinguish the Vps8Dp-and Dop1p-defined zones of the bladder. We exposed the dual-labeled cells described above to mild osmotic stress for a brief period, as described in Materials and Methods, and then measured the same radial plots for the pairs of CV proteins.

Interestingly, the distance between the peak intensities of Dop1p and Vps8Dp increased upon osmotic stress (Fig. 5I and 5J) while that between Vps8Dp and Vma4p decreased (Fig. 5K and 5L). The change in the distribution of Vma4p upon osmotic shock is driven primarily by a striking response to the osmotic stress, namely what appears to be the contraction or collapse of the spongiome (Fig. 5K right panel). As a result of this contraction, Vma4p signal becomes much more heavily concentrated near the bladder periphery. A composite model including the distributions of Dop1p, Vps8Dp, and Vma4p within the CVC is shown in Figure 5M.

### 7. CVC-localized Vps8Dp and Dop1p exchange with large cytosolic pools

That the CVC is a highly dynamic structure is apparent both from its periodic contraction as well as the tubule extension and branching visualized above. Numerous Rabs and SNAREs localize to the CVC in ciliates, suggesting that the mechanisms underlying CVC dynamics involve active vesicle budding and fusion (Plattner, 2013, Plattner, 2010, Bright et al., 2010, Turkewitz and Bright, 2011, Schonemann et al., 2013). Similarly, Vps8Dp, as part of CORVET, is likely to be directly involved in vesicle trafficking.

We asked whether Vps8Dp, together with other CVC proteins identified in this study, are dynamically associated with CVC membranes. We performed Fluorescence Recovery after Photobleaching (FRAP) by briefly bleaching a defined sector of the CVC in immobilized cells and measuring the time of fluorescence recovery, which depends on exchange with fluorescent proteins from outside of the bleached area. A limitation of these experiments is that we could be certain of bleaching the entire CV but not the spongiome, given the expanse of the latter. These experiments were done using a confocal microscope, so the appearance of the fluorescent signals differs for some tagged proteins compared to the epifluorescence imaging shown in earlier figures.

Figure 6A-F show FRAP results for Dop1p and Vps8Dp, both of which are peripheral membrane proteins. The data reveal both proteins dynamically exchange with pools in the non-bleached area, but that exchange of Vps8Dp appears more rapid than that of Dop1p (Fig. 6B, 6E, S6 and S7). The extent of the recovery of each protein after bleaching may be limited by the relative sizes of the CV-associated vs cytosolic pools. To gain some insight into this, we used these images to estimate the relative amounts of each protein pool at steady state that is localized to the CV vs non-CV area, subject to the limitation regarding defining the precise spongiome borders. This analysis suggests that both Dop1p and Vps8D include significant pools of cytosolic protein, but that this fraction is higher for Vps8Dp (Fig. 6C, 6F, S8 and S9).

**Figure 6:**
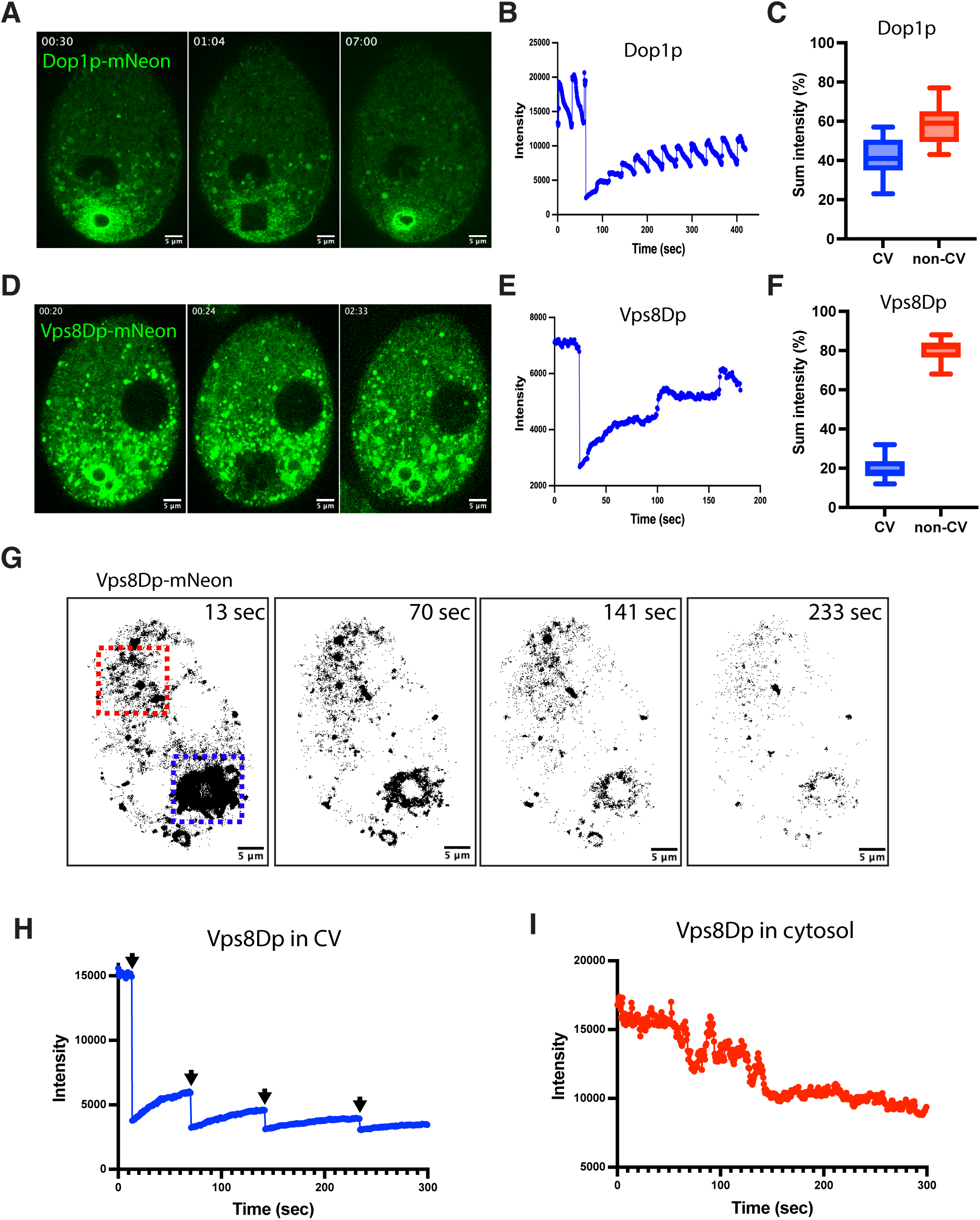
CV-localized Dop1p and Vps8Dp differentially exchange with large cytoplasmic pools. A: Cell expressing Dop1p-mNeon analyzed by FRAP. Growing cells were immobilized and live imaged with a Marianas spinning disc confocal microscope with FRAP tool. Three images were extracted from a video (Movie 12) showing a photobleaching event and recovery. Left panel: cell before photobleaching at 30s. Middle panel: cell immediately after photobleaching a region containing the CV and periphery at t=1:04. The bleached area appears as a dark square. Right panel: cell at t=7:00. B: The recovery after photobleaching data were analyzed by FIJI image process software (see details in Materials and Methods). The fluorescence intensity in the photobleached area was plotted using GraphPad Software Prism. The spikiness of the trace is due to periodic contraction of the CV. C: The percentage of Dop1p in the CV and non-CV areas. The borders of the areas for quantification in 17 independent images (Figure S5) were determined by using FIJI software and the sum intensities of the designated areas were measured, as described in Materials and Methods. D: Cell expressing Vps8Dp-mNeon analyzed by FRAP. Three images were extracted from a video (Movie 13) showing a photobleaching event and recovery, with panels as in Fig. 8A but with the bleaching event at t=0.24 and recovery at t=2:33. E: Recovery after photobleaching, plotted as in Fig. 6B. F: The percentage of Vps8Dp in the CV versus non-CV areas. The approaches were identical to those used in Figure 6C and the images and data are shown in Figure S6 and Table S3. The experimental procedures and analytical methods used in Figures 6D-F are identical to those used in Figure 6A-C, respectively. G: Cell expressing Vps8Dp-mNeon analyzed by FLIP. Four images were extracted from a video (Movie 14) in which the CV (bottom right, dotted blue square) was bleached four times successively at t=13s, 70s, 141s and 233s. The fluorescence intensity was measured both in the dotted blue square and in an anterior zone of the cytoplasm (dotted red square). The fluorescence intensities in both zones were analyzed by FIJI image process software. H: The fluorescent signal of CV-localized Vps8Dp-mNeon (in the dotted blue square) decreased and recovered repeatedly after bleaching. Arrows indicate the four photobleaching events. I: The anterior cytoplasmic Vps8dp-mNeon signal (in the red dotted square) decreased after photobleaching the CV.

To ask more directly whether the cytosolic pool of Vps8Dp exchanges with the CVC pool, we used FLIP (Fluorescence Loss in Photobleaching). We repeatedly bleached the CVC of immobilized cells expressing Vps8Dp-mNeon over a period of minutes, while monitoring the Vps8Dp-mNeon fluorescence at a distant site in the cell anterior (Fig. 6G). The results are consistent with exchange between cytoplasmic and CV-localized pools, since Vps8D fluorescence in the cell anterior decreases upon bleaching of the CV pool (Fig. 6H and 6I).

### 8. Vma4p and Dop1p undergo exchange via different mechanisms

Vma4p, unlike Dop1p and Vps8Dp, is a subunit of an integral membrane protein complex. In addition, unlike Dop1p and Vps8Dp, the entire pool of detectible Vma4p is localized to the CVC. We used FRAP to ask whether Vma4p is mobile within the spongiome membrane (Fig. 7A, 7B and S10). After bleaching a subsection of the spongiome, we observed that Vma4p-mNeon fluorescence initially begins to recover at one edge and then progressively spreads (Fig. 7C). This is consistent with the idea that Vma4p is mobile within the plane of the membrane, and that exchange is primarily due to diffusion along the tubules. In contrast, when the same analysis was performed on cells expressing Dop1p-mNeon, the fluorescence recovery was not directional but instead occurred in patches throughout the bleached area (Fig. 7D and S6A). This result is consistent with the idea that Dop1p exchange occurs via a cytosolic pool, which could be either soluble or vesicle-associated. This difference between the spatial patterns of fluorescence recovery of Vma4p and Dop1p could also be seen in kymographs plotting the recovery of fluorescence (Fig. 7E).

**Figure 7:**
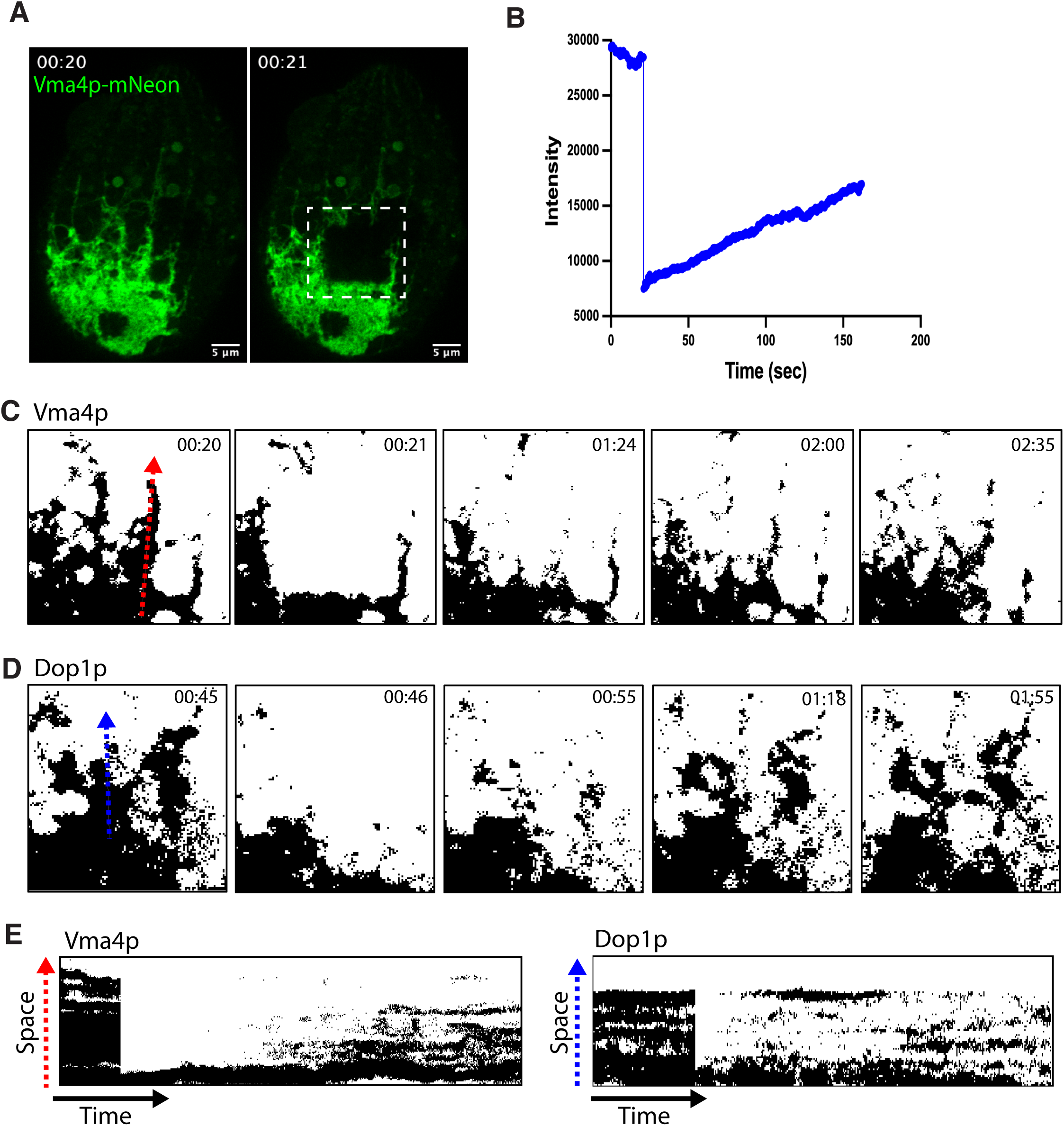
Vma4p and Dop1p exchange via different mechanisms. A: Cell expressing Vma4p-mNeon analyzed by FRAP. Two images were extracted from a video (Movie 15) to demonstrate a FRAP experiment in which a sector of the reticulum was bleached. Left panel: cell before photobleaching at t=20s. Right panel: bleaching event at t=21s, resulting in a dark square. Growing cells were immobilized and live imaged with a Marianas spinning disc confocal microscope with FRAP tool. The data were analyzed by FIJI image process software. B: Recovery after photobleaching. The fluorescent intensity in the indicated area before and after photobleaching was plotted using GraphPad Software Prism. C, D and E: Vma4p recovery after photobleaching includes diffusion within the tubule membranes, while Dop1p exchanges with a soluble pool. C: Magnified image of region indicated in Fig 7A right panel. Five images were extracted from a video (Movie 15) to illustrate that the recovery of Vma4p-mNeon fluorescence is directional and proceeds from the bottom of the field. This is also shown in the kymograph (Figure 7E), based on tracking signal intensity/time along the red arrow shown in Fig 7C. To illustrate the recovery process of Dop1p in the tubules, five images are shown (Figure 7D) and a kymograph (Figure 7E) from spongiome-localized Dop1p FRAP video footage (Figure S5A and Movie 16). The red/blue dotted lines indicate the space used to trace the change of signal intensities over time for kymograph analysis, by FIJI image process software.

### 9. The CVC is isolated from bulk endocytic traffic

Organelles in the endomembrane network are maintained by a balance of inward and outward membrane traffic. The relevant trafficking pathways for the CVC are poorly understood in any organism. Recent analysis in *Dictyostelium* points to the involvement of endosomal fusion machinery, though the specific roles are as yet unexplored (Manna et al., 2023). Similarly, an important role for endosomal trafficking to the Ciliate CVC is consistent with the markers we have established above, since CORVET in other organisms functions as an endosomal or endolysosomal tether-fusion complex, and the DOPEY family proteins are similarly endosome-associated. Both vacuolar ATPases and lipid scramblases function in many compartments including endosomes (Marshansky and Futai, 2008, Hankins et al., 2015).

In *Tetrahymena* as in many other cells, fluorescent styryl dyes can intercalate into the outer leaflet of the plasma membrane and be taken up into endocytic vesicles. Such dyes can therefore be used to trace endocytic trafficking (Betz and Bewick, 1992, Betz et al., 1996, Elde et al., 2005). We briefly incubated cells expressing tagged CV markers with red fluorescent FM4-64, and looked for overlap between the endocytosed FM dye and either Dop1p (Fig. 8A) or Vma4p (Fig. 8B). Strikingly, there was no detectible overlap after 15 min or 30 min of FM update, with the same result after overnight incubation (Fig. S11). Thus, the CVC in Tetrahymena appears to be isolated from bulk endocytic traffic.

**Figure 8:**
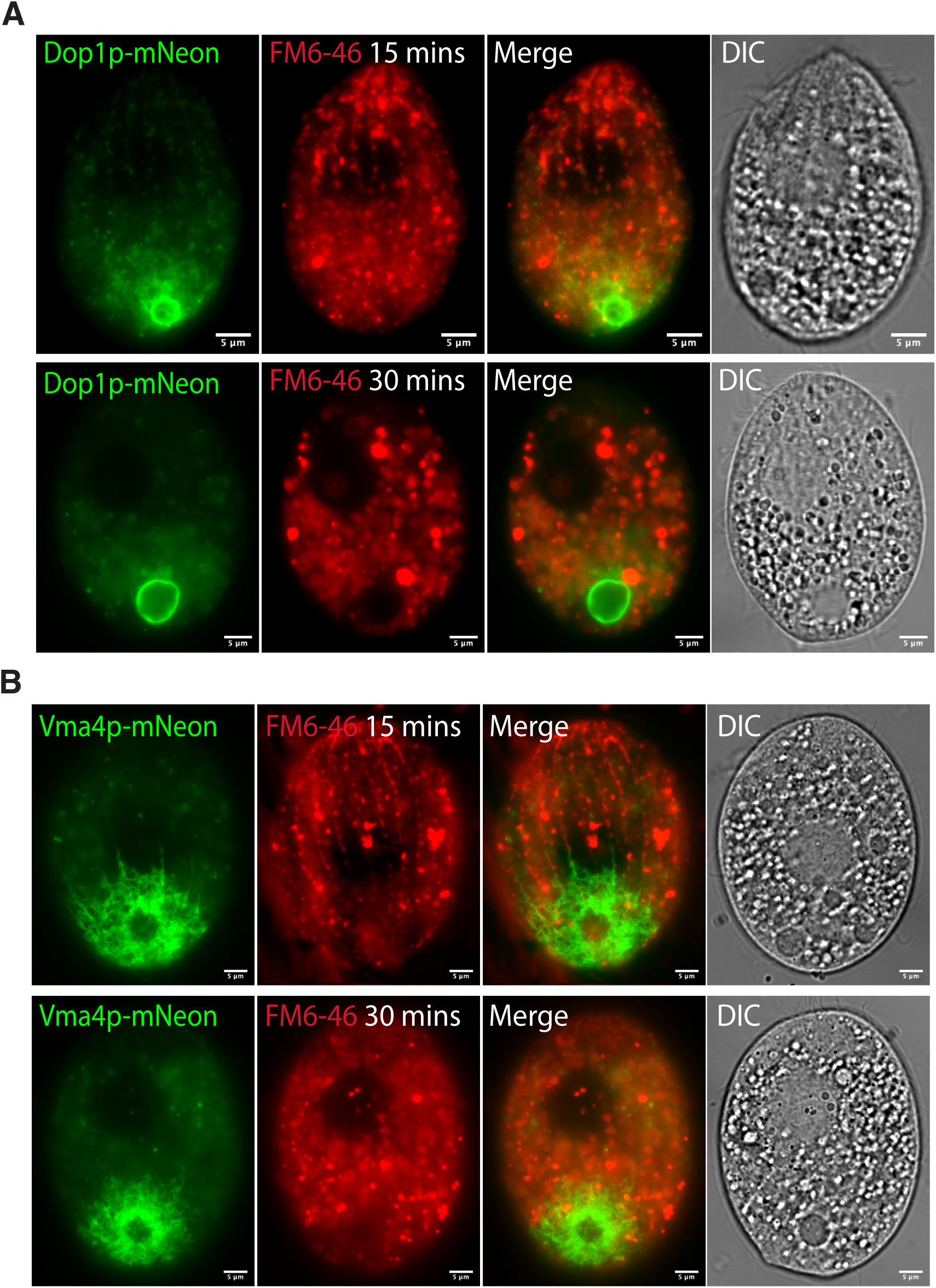
The CVC does not receive bulk endocytic traffic. A: Cells expressing Dop1p-mNeon were incubated with the red fluorescent endocytic tracer FM4-64 for 15m (top panel) or 30m (bottom panel). B: Cells expressing Vma4p-mNeon were incubated with FM4-64 for 15m (top panel) or 30m (bottom panel). There was no significant overlap between FM4-64 labeling and either of the CVC protein markers. Growing cells were incubated with FM4-64 as described in Materials and Methods, immobilized, and live imaged with a Zeiss Axio Observer 7 system.

## Discussion

Though the contractile vacuole has been recognized by biologists for more than three centuries, the structure and particularly the mechanisms underlying periodic contraction are still largely obscure. This lack of knowledge is partly attributable to the fact that while CVs are widely distributed among eukaryotes, they are not present in fungi, animals or plants where most cell biology has been pursued. A corollary hurdle is that many of the organisms that do bear CVs have been inaccessible to many genetic and molecular approaches critical to analyze such complex cell biological structures. CVs are prominent in both marine and fresh-water ciliates, and experiments and observations in Paramecium sp. have been important in generating our current understanding of the CV (McKanna, 1976, Kitching, 1939). Among the Ciliates, the species with the best developed toolbox is *Tetrahymena thermophila* (Ruehle et al., 2016). In this manuscript, we establish a set of live markers for the CV in this organism. While some cv-associated proteins have previously been identified in both Tetrahymena and Paramecium sp., the uniqueness of the localization has rarely been established, and/or other limitations precluded their use as live markers. We previously reported a set of Rab GTPases which, as over-expressed GFP-tagged proteins, appeared highly concentrated at the CV (Bright et al., 2010). However, these tagged proteins may not be readily visible when expressed at endogenous levels, a problem we have encountered with many Rabs in this organism (unpublished).

As shown in this manuscript, four proteins, two of which are peripherally associated with the membrane and two which are membrane-embedded or subunits of integral membrane complexes, can be tagged and used to visualize CVC dynamics when expressed at endogenous levels. These results significantly expand our ability to analyze the CVC in ciliates. While the bladder could already be visualized even in the 18^th^ century by light microscopy, our ability to capture videos of fluorescent bladders has revealed new details of the contractile cycle. The observation that Dop1p localized to a subtly different position at the bladder periphery compared to Vps8Dp and Scr7p, and that these zones show differential responses to osmotic challenge, should allow the mapping of subdomains in this complex organelle. In Paramecium, where the CVC is morphologically more complex than in Tetrahymena, there is a notable substructure within the spongiome where distal versus peripheral tubules show differential occupancy by vacuolar ATPase as well as an aquaporin (Ishida et al., 2021).

Fluorescently-tagged proteins that label the CVC structures also permitted us to recognize novel features of CVC duplication in mitotic cells, including the discovery that newly forming Dop1p-labeled bladders appear during early stages of cell division, as two rings intermediate in size between pores and mature bladders. Previous studies of CVC duplication have relied on the pores, visible using silver staining, and these appeared only at relatively later stages of cell division (Frankel, 1992, Frankel, 2000, Cole and Gaertig, 2022). Our observations thus advance the timeframe for a key morphogenetic event during cell division. The identification of CVC-specific genes and proteins now presents opportunities for exploring developmental regulation, extending recent studies on cortical determinants for position-specific organelle biogenesis in ciliates. These studies have highlighted the crucial role of morphogen gradients mediated by signaling molecules in defining cortical pattern formation during cell division (Jiang et al., 2017, Jiang et al., 2019a, Jiang et al., 2019b, Jiang et al., 2020). One open question is whether CVC-specific proteins interact with the newly identified morphogens to influence the positioning and assembly of the new CVC. The precise positioning of new CV pores has been proposed to be regulated by their positioning relative to other organelles, rather than being fixed to a specific cortical pattern (Ng, 1977, Ng and Frankel, 1977). Intriguingly, we found that tubules labeled with Dop1p puncta as well as Vma4p, extend along meridians during cell division and potentially establish a connection between the pre-existing and new CVCs, so that the latter may derive in part from the former either informationally and/or compositionally.

The mechanisms underlying bladder contraction are unknown. CV bladders that were micro-surgically excised from *Paramecium* briefly continued to undergo periodic contraction, suggesting that remodeling does not depend on an organized cytoskeleton (Tominaga et al., 1998a, Tani et al., 2000, Tani et al., 2002). Ultrastructural studies have revealed what may be an unusual packing of CV membranes during contraction, with possible similarity to lipid cubic phases (McKanna, 1976, Tominaga et al., 1999, Allen, 2000). A distinct lipid composition of the *Dictyostelium* bladder has been proposed to directly drive contractile cycles in a model that invokes flipping across the membrane bilayer (Heuser et al., 1993), but as yet this model is untested. Our finding that the predicted lipid scramblase Scr7p is strongly concentrated at the CVC corroborates that lipid transfer plays a role in this organelle, which we hope to elucidate in the future. We found that Scr7p closely co-localizes with Vps8Dp, and distal to the localization of Dop1p at the bladder periphery, but higher resolution imaging will be needed to understand precisely what structures are involved.

The homologs of *DOP1* and *VPS8D* in other organisms are endosomal proteins (Balderhaar and Ungermann, 2013, Moliere et al., 2022a, van der Beek et al., 2019a). An inference, consistent with the implications from work in other organisms, is that the CVC is a specialized endosomal compartment. However, the CVC in Tetrahymena does not appear to directly receive traffic from endocytic vesicles, judging from our results with the endocytic tracer FM4-64. Those results also suggest that there is little or no diffusion in the plane of the membrane through the CV pore, consistent with results in Amoeba proteus (Nishihara et al., 2007). Similar results have been reported in *Dicytostelium*, though in that case FM dye-labeling of the CVC was seen in stressed cells (Heuser et al., 1993, Gerisch et al., 2002). A better understanding of ways in which the CVC is integrated into the endomembrane network will be facilitated by identifying and analyzing additional CVC-specific proteins in experimentally tractable organisms. In parallel, developing tools to manipulate a wider phylogenetic range of eukaryotes should make it possible to determine whether CVs in diverse organisms depend on evolutionarily shared mechanisms, and therefore to shed insight into the origins of this organelle (Manna et al., 2023).

## Materials and Methods

### Cell strains and culture conditions

*Tetrahymena thermophila* strains used in this work are listed in Table S1 with culture conditions as previously described (Gorovsky et al., 1975). Cells were grown in SPP medium (2% proteose peptone (GIBCO, 211684), 0.1% yeast extract (BD, 212750), 0.2% dextrose (ACROS, 41095-5000) and 0.003% ETHYLENEDIAMINE-TETRAACETIC ACID, Ferric-Sodium Salt (SIGMA)) supplemented with 100 μg ml^-1^ Normocin^TM^ (InvivoGen), an antimicrobial reagent. Cells were grown at 30°C with agitation at 100 rpm. Cell densities were measured using a spectrophotometer (Thermo Spectronic Unicam), where OD550 = 1 corresponds to ∼ 1 *×* 10^6^ cells ml ^-1^ (David L. Spector, 1998) or a Z1 Coulter Counter (Beckman Coulter Inc., Indianapolis, Indiana). Cultures were generally used during the log phase of cell growth (cell density 2-4 × 10^5^ cells ml^-1^).

### Biolistic Transformation

50 ml *Tetrahymena* cultures were grown to 500,000 cells ml^-1^ and starved for 10–16 h in 10 mM Tris-HCl buffer, pH 7.4, at 30°C with agitation. Biolistic transformations were performed as described previously (Sparvoli et al., 2018, Cassidy-Hanley et al., 1997). Transformants were identified after 3 days selection in paromomycin sulfate (120 μg ml^-1^ with 1 μg ml^-1^ of CdCl2), and then serially passaged 5×/week for ∼4 weeks in decreasing concentrations of CdCl2 and increasing concentrations of paromomycin sulfate. Reagents were from Sigma-Aldrich unless otherwise noted.

### Generation of *VMA4* knockout strains

The Macronuclear open reading frame (ORF) of the *VMA4* gene (TTHERM_00193660) was replaced with the paromomycin (Neo4) drug resistance cassette (Mochizuki, 2008) via homologous recombination with the linearized vectors pVMA4KO-Neo4. PCR was used to amplify the fragments of the genomic regions upstream (5′UTR), 1248 bp and downstream (3′UTR), 1481 bp of the ORF. The amplified fragments were subsequently cloned into restriction sites generated by NotI/PstI for the upstream fragment and BamHI/XhoI for the downstream fragment, flanking the Neo4 cassette of the pNeo4 vector, by Quick Ligation (New England, Biolabs Inc.) The primers used to create these constructs are listed in Table S2. The construct was linearized by digestion with SacI and KpnI and transformed into CU428.1 cells by biolistic transformation. Restriction enzyme reagents were from NEW ENGLAND BioLabs.

### Endogenous expression of the CVC-related genes with mNeon fluorescent tags

The p2mNeon-6myc-Neo4 vector (Sparvoli et al., 2018) was designed to integrate two mNeonGreen fluorescent tags at the C-terminus of a targeted macronuclear target gene. For each target gene, we used PCR with a proofreading Taq polymerase (Roche, Expand Long Template PCR System) to amplify the C-terminal region (∼0.5-1 kb, but not including the stop codon) as a 5’-flanking fragment, and the 3’UTR as a 3’-flanking fragment. The PCR products were verified by agarose gel electrophoresis and the purified DNA fragments were cloned into p2mNeon-6myc-Neo4 vector at the Xbal site for 5’-flanking fragments and the Xhol site for 3’-flanking fragments, respectively, by using NEBuilder HiFi DNA Assembly. The final construct was verified by restriction mapping, and constructs were isolated from 50 ml bacterial cultures using a plasmid DNA extraction kit (Takara Bio USA, Inc. NucleoSpin Plasmid). DNA concentrations were measured by a NanoDrop spectrophotometer. 20 µg of each construct was linearized with PvuII and then purified for each biolistic transformation. The primers used are listed in Table S2.

### Dual endogenous gene tagging

The construct of pVPS8D-3xmCherry-2HA-CHX was previously constructed to endogenously express mCherry-tagged Vps8Dp (Sparvoli et al., 2020). We modified the construct to replace the cycloheximide (CHX) selection cassette (bounded by PstI and XhoI sites) with a cassette conferring resistance to puromycin, which we obtained by amplifying the selectable Pur4 marker by PCR from the pPur4-opt vector (Iwamoto et al., 2014). Assembly was using the NEBuilder HiFi DNA Assembly method. The pVPS8D-3xmCherry-2HA-PUR4 was biolistically transformed into cells already expressing Dop1p-mNeon or Vma4p-mNeon.

### Cell immobilization for live imaging

We used two methods for cell immobilization for live cell imaging. For both methods, we prepared a single slide at a time and viewed it immediately. The first method relied on the pressure exerted on small volume samples under a cover slip. Cells were first concentrated by centrifugation to 2-5 x 10^6^ cells ml^-1^. 6 μl of sample was applied to the slide, and immediately overlayed with a 22 × 22-1 cover slip. The resulting pressure frequently results in cell immobilization, which could be verified by monitoring ciliary beating on the cell surface at the same time that cell viability could be monitored by observing the periodic contraction of the contractile vacuole, both in the DIC channel. For the second method, cells were pelleted and resuspended in a thermoreversible gel on ice, whose viscosity increases when it is warmed to room temperature. Cells at the same high density were mixed well with CyGEL^TM^ (ab109204, ABCAM) (the mix ratio optimized for each experiment) and immediately mounted with a cover slip. This approach results in many cells being effectively immobilized in the thin gel. For both methods, we carefully watched for proliferation of large cytosolic vesicles. Growing cells have large food vacuoles derived from the oral apparatus. No new food vacuoles form in immobilized cells, so any substantial increase in the number of such vesicles is likely to represent autophagosome formation as part of a stress response. We rejected any cells showing such an increase.

### Cell fixation for imaging

Cells were fixed in a final concentration of 4% paraformaldehyde in 10 mM Tris-HCl buffer, pH 7.4 (stock: 16% paraformaldehyde solution in distilled water, EM grade, 15710, Electron Microscopy Science) for 10-30 minutes at room temperature. They were then washed repeatedly in PBS to reduce the background autofluorescence, before staining with DAPI (4′,6-diamidino-2-phenylindole) at final concentration 50 ng ml^-1^ for 10 minutes for nuclear staining, and/or staining with antibodies for immunofluorescence. To improve preservation of the tubular structure of the spongiome, we fixed cells with 3% paraformaldehyde for 10 minutes on ice. Images were captured by using a Carl Zeiss Microscope stand Axio Observer 7 system or Marianas Yokogawa-type spinning disc inverted confocal microscope.

### Fluorescence Recovery after Photobleaching (FRAP)

Cells were immobilized as described above. FRAP experiments were performed using a Marianas Yokogawa-type spinning disc inverted confocal microscope with a 100x/NA1.45 oil (Alpha Plan-Fluar) objective. The system featured fast shutter speeds and channel switching for high-speed imaging and a vector high-speed point scanner for bleaching, controlled using Slidebook software. For each experiment, we first identified a well-immobilized cell and verified the fluorescence signal with a ∼100-500 ms exposure. We then used the time-lapse mode to capture images with the fastest camera speed for a pre-photobleaching record and then used the screen selection tool and real-time imaging to draw a ROI (region of interest) for photobleaching.

### Labeling endocytic compartments with FM4-64

Cell cultures were incubated for 15 min, 30 min or overnight with 5 μM FM4-64 (Invitrogen).

### Image analysis

The image processing package FIJI was used for image processing and analysis (Schindelin et al., 2012). Processing of raw images included: image cropping and rotation; bleach correction; background subtraction; adjustment of brightness/contrast; color switching; selection of images from z-stacks; intensity threshold adjustments; measurements of parameters including areas, mean gray values, and integrated densities. The Radial Profile plugin was used to measure and plot the intensity of fluorescence signal along specified radii. A region of interest (ROI) corresponding to the CV bladder was created as a circle whose center corresponded with the apparent bladder center and whose circumference included the entire visible CV periphery. The normalized integrated intensity of the tagged fluorescent protein was calculated along many radii, and the average intensity at each radial position was plotted as a function of distance from the circle center.

To quantify fluorescence intensities in the CV vs cytosol, all images were captured using the same microscopy settings. The raw images were used to adjust the threshold signal intensity for segmenting the whole cell area from extracellular background, and for segmenting the CV area from the cytosol, both using Binary selection with Fill Holes. The sum fluorescence intensities of the whole cell and the CV area were then measured. For FRAP analysis, the time-lapse images were separately adjusted by bleach correction for the pre-photobleaching series and after-photobleaching series. The intensity thresholds were adjusted for background subtraction, and intensity changes over time were measured in ROIs. Intensities were plotted using GraphPad Software Prism.

### Osmotic challenge

Cells expressing tagged proteins were grown to log growth phase in SPP medium. For hypo-osmotic treatment, cells were pelleted by centrifugation for 1 minute and resuspended gently in 10 mM Tris-HCl buffer, pH 7.4 for 1 minute. This process was repeated and the cells were then incubated in the same buffer, at room temperature, for 10 minutes before being fixed for imaging.

### Immunofluorescence analysis

Cells (2×10^5^) were fixed with 4% paraformaldehyde for 30 minutes at room temperature, and immunolabeled as previously described (Briguglio et al., 2013, Bowman and Turkewitz, 2001). Briefly, cells were first incubated for blocking with 5% BSA (bovine serum albumin, BP1600-1 Fisher Scientific) in PBT (0.3% Triton X-100 (ACROS) in PBS) for one hour at room temperature and then incubated with the primary antibodies in the same blocking buffer. Anti-α-Tubulin antibody staining was at 1:2000 (clone DM1A ZooMAb® Mouse Monoclonal, Sigma ZMS1039) for overnight at 4°C, followed by three washes in 0.3% Triton X-100 in PBS. The secondary antibodies were Texas Red-conjugated anti-mouse IgG, incubated for one hour at room temperature (1:1000, Texas Red Goat anti-Mouse IgG, Invitrogen T2767), and similarly washed. Cell nuclei were stained with 4′,6-diamidino-2-phenylindole (DAPI) (50 ng ml^-1^). Microscopic slide mounting and sealing for microscopy imaging.

### Expansion microscopy

The method was adapted from (Damstra et al., 2022). Growing cells were fixed with 4% PFA for 10 min at room temperature. Fixed cells were incubated with 0.1 mg ml-1 acryloyl X-SE (AcX, 2541676, Invitrogen) for a minimum of 3 hours in a dark humidified chamber. For gelation, 194 µl (for 2 samples) of the monomer solution (sodium acrylate 1.1M; acrylamide 2 M; N,N’-methylenebisacrylamide 0.009% in PBS) was prepared and kept on ice. Tetramethylethylendiamine (TEMED, 11042F, GIBCO BRL, 3 µl of 10% stock in water) and ammonium persulfate (A3678, SIGMA, 3 µl of 10% stock in water) were added and quickly mixed on ice, and 97 µl of gelation solution was mixed on the slide with 3 µl of concentrated fixed cells. The slides were incubated in the dark at 37°C for 1 hour, after which proteinase K (42-700, Apex Bioresearch) was added (final concentration 7.5 units ml-1) for at least 2 hours in the dark at 37°C. Afterward, the gels were gently removed and transferred to 15 cm petri dish with three successive water washes water for promote expansion. The extent of anisotropic expansion was estimated by measuring the diameter of the gels in two orthogonal directions. For imaging, small pieces were cut from the gel and mounted (cells facing down) on the coverslip (Glass Bottom Microwell Dishes, Part No.: P35G-1.5-10-C) which had been pre-coated by adding ∼70 µl of 20% poly-L solution and dried for 30-60 minutes on a 37°C heat block. Finally, 20 µl of water was added and the sample covered with a 2nd cover slip.

## Acknowledgements

We thank Daniela Sparvoli for the Vps8D-mCherry construct, and Masaaki Iwamoto (Nihon University) for the puromycin drug selection plasmid. At the University of Chicago, we thank Vytas Bindokas and Christine Labno for help at the Integrated Light Microscopy Core Facility. In the Turkewitz laboratory, we thank Patrick Jiang, Maya Waarts, and Josefina Hernandez for helpful discussions.

