## Supplementary figures and images for "Structure and dynamics of the contractile vacuole complex in *Tetrahymena thermophila*"

### Fig. S1

Fig. S1

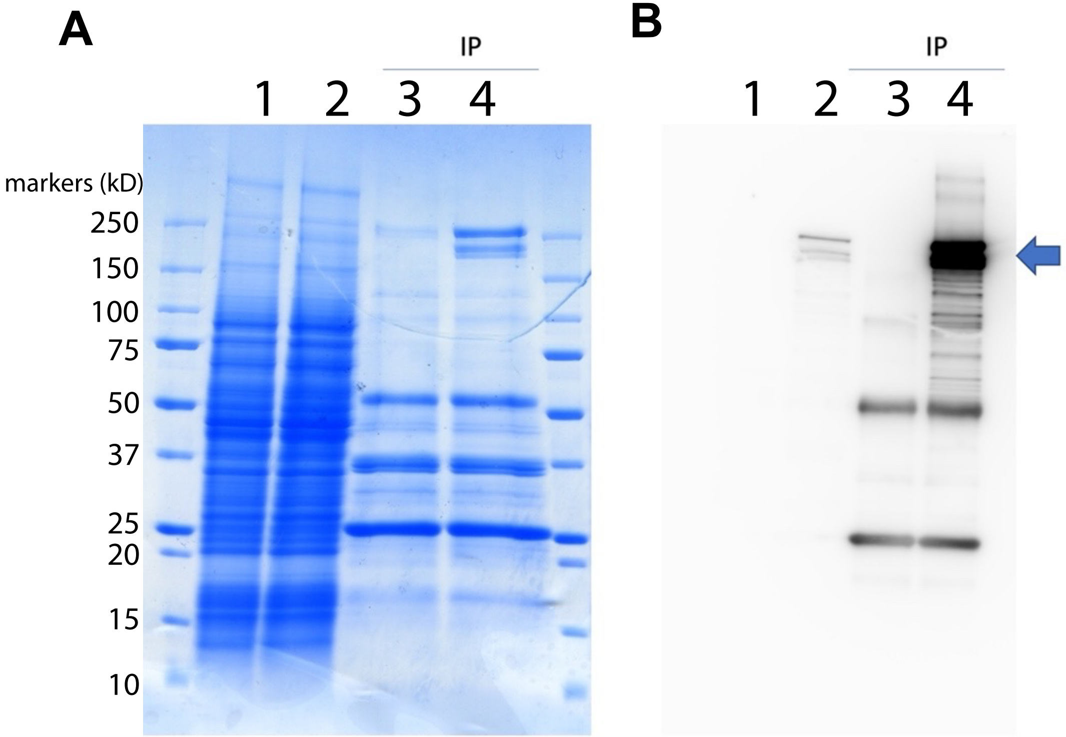

### Fig. S3

Fig. S3

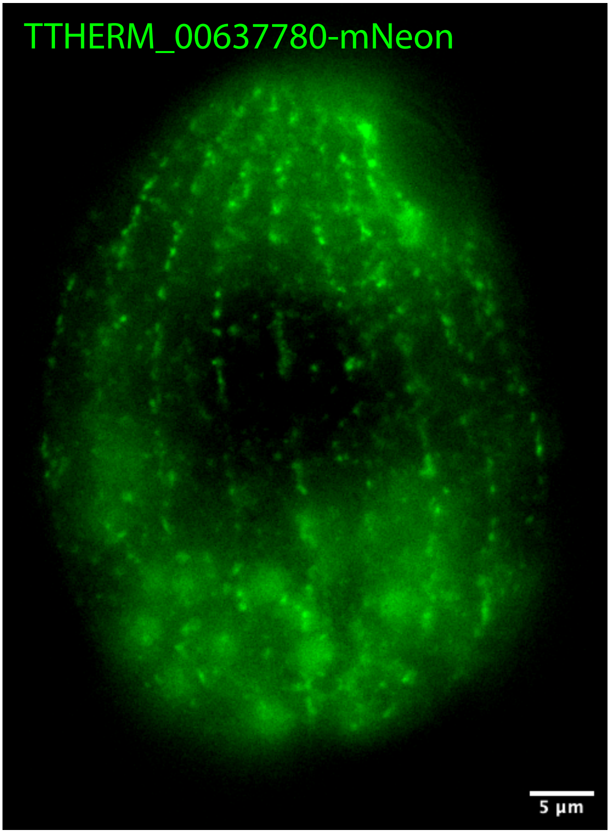

### Fig. S4

Fig. S4

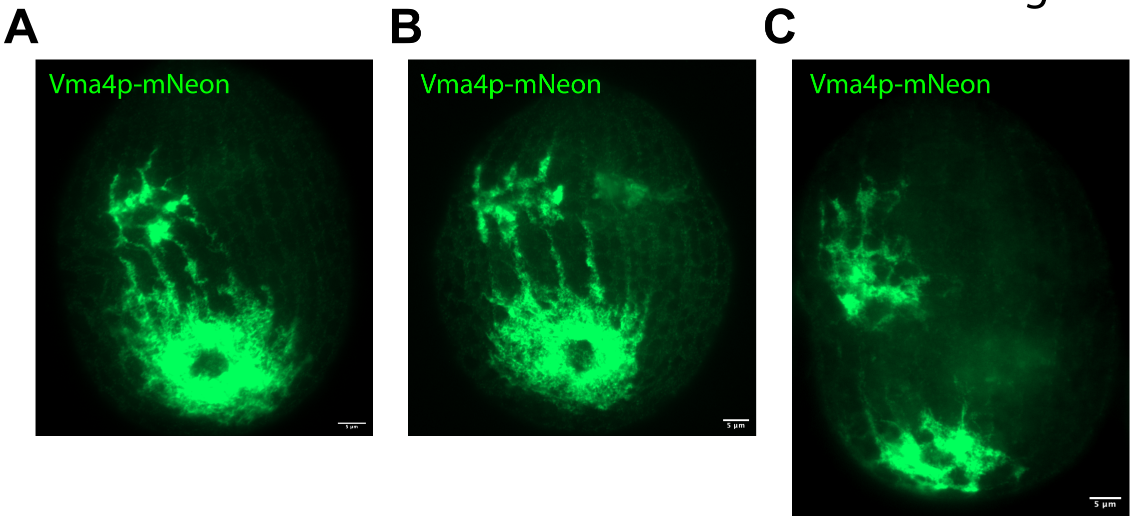

### Fig. S5

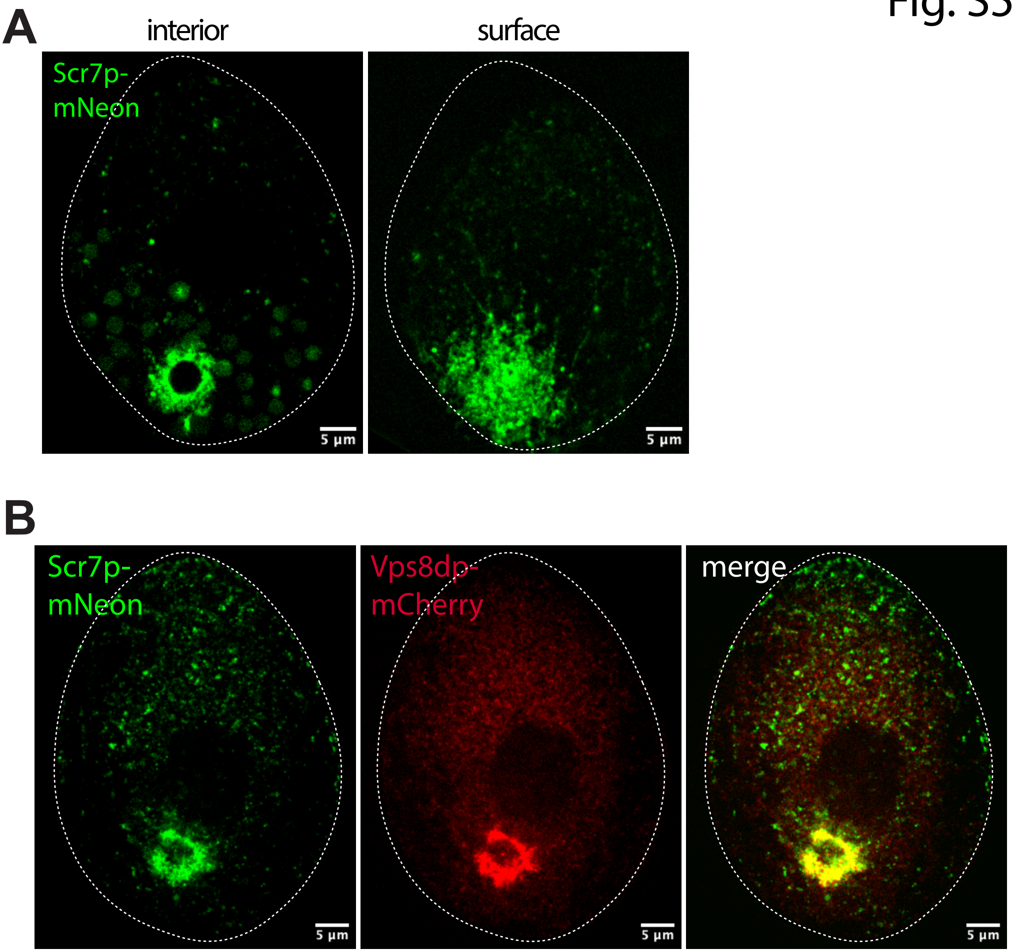

### Fig. S6

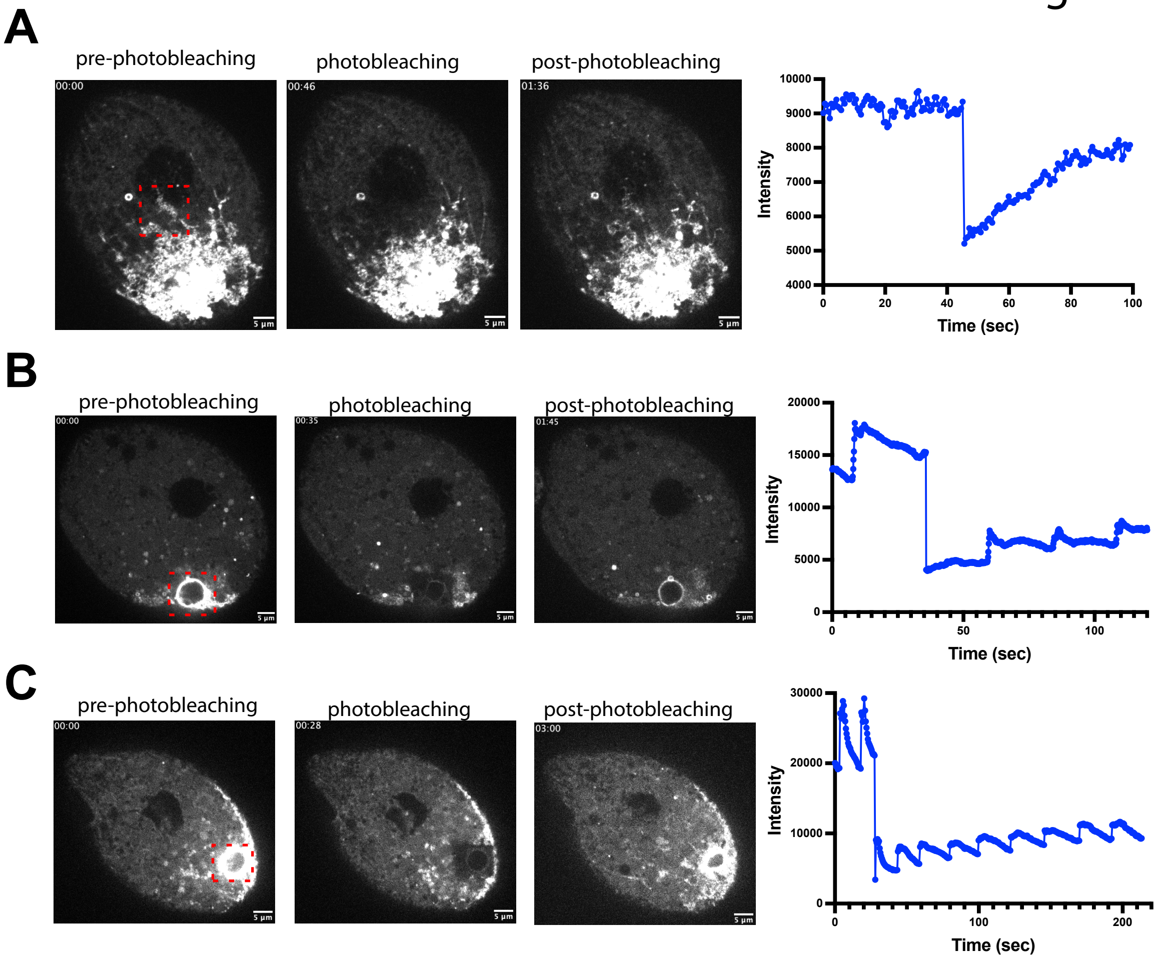

### Fig. S7

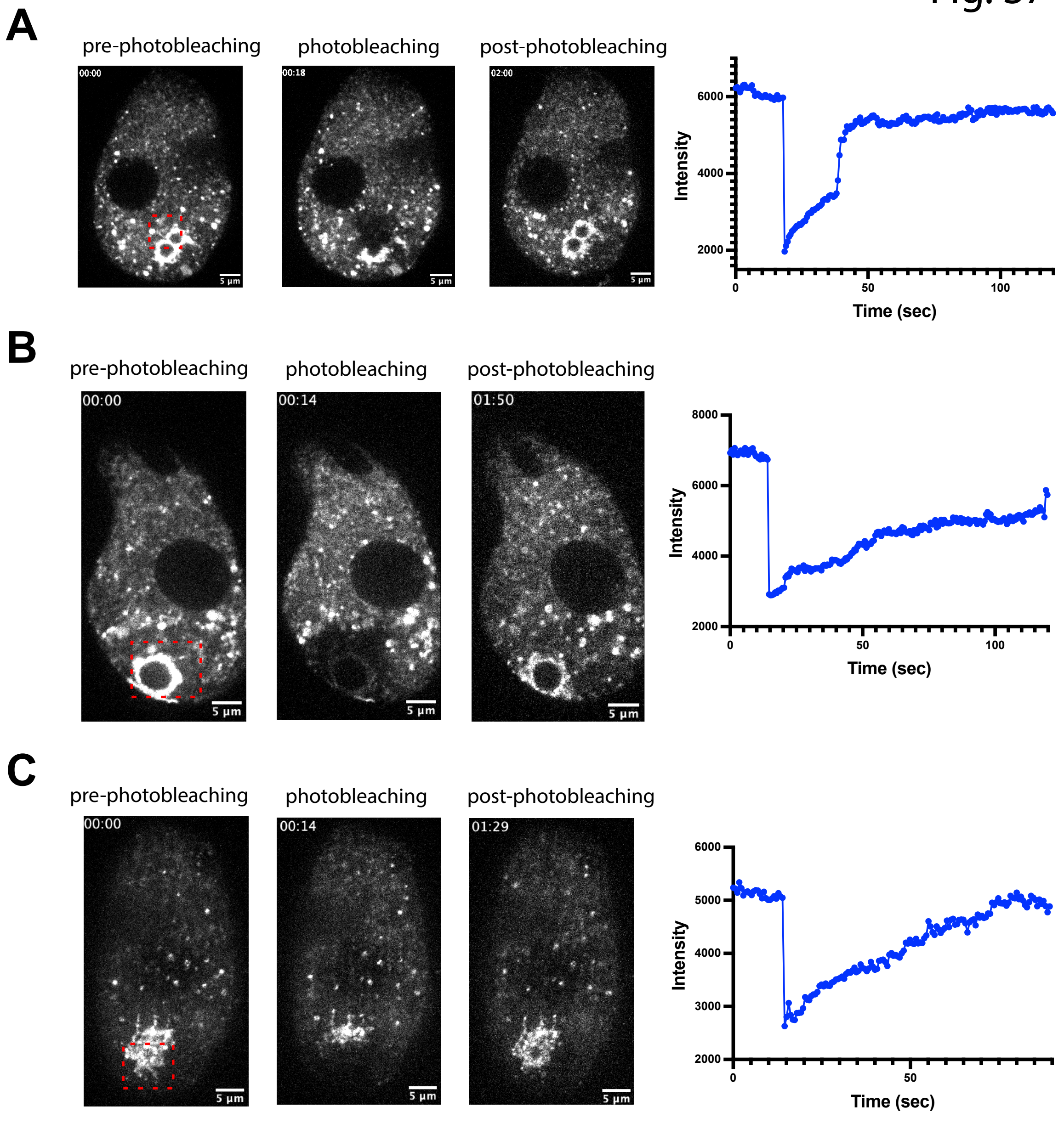

### Fig. S8

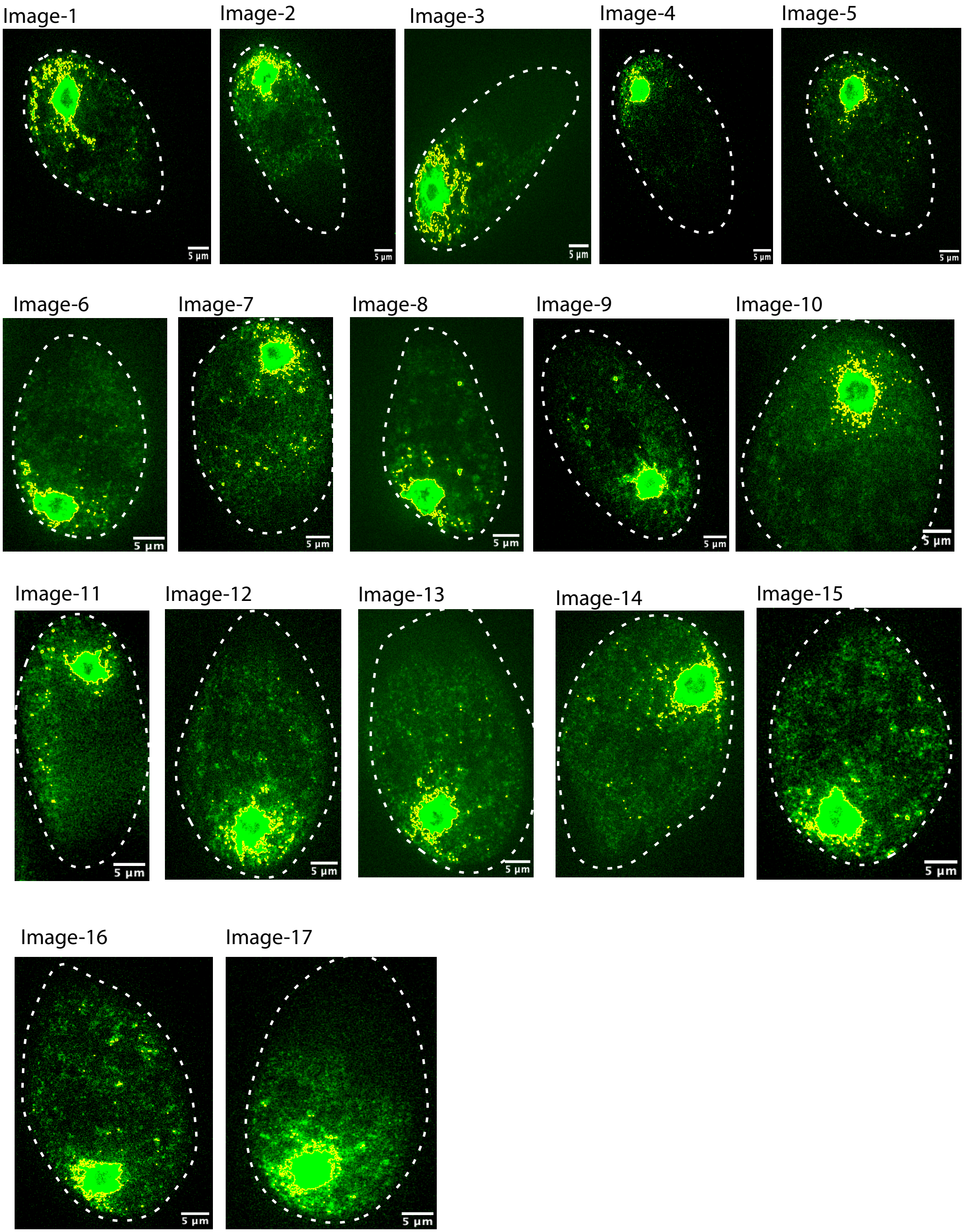

### Fig. S9

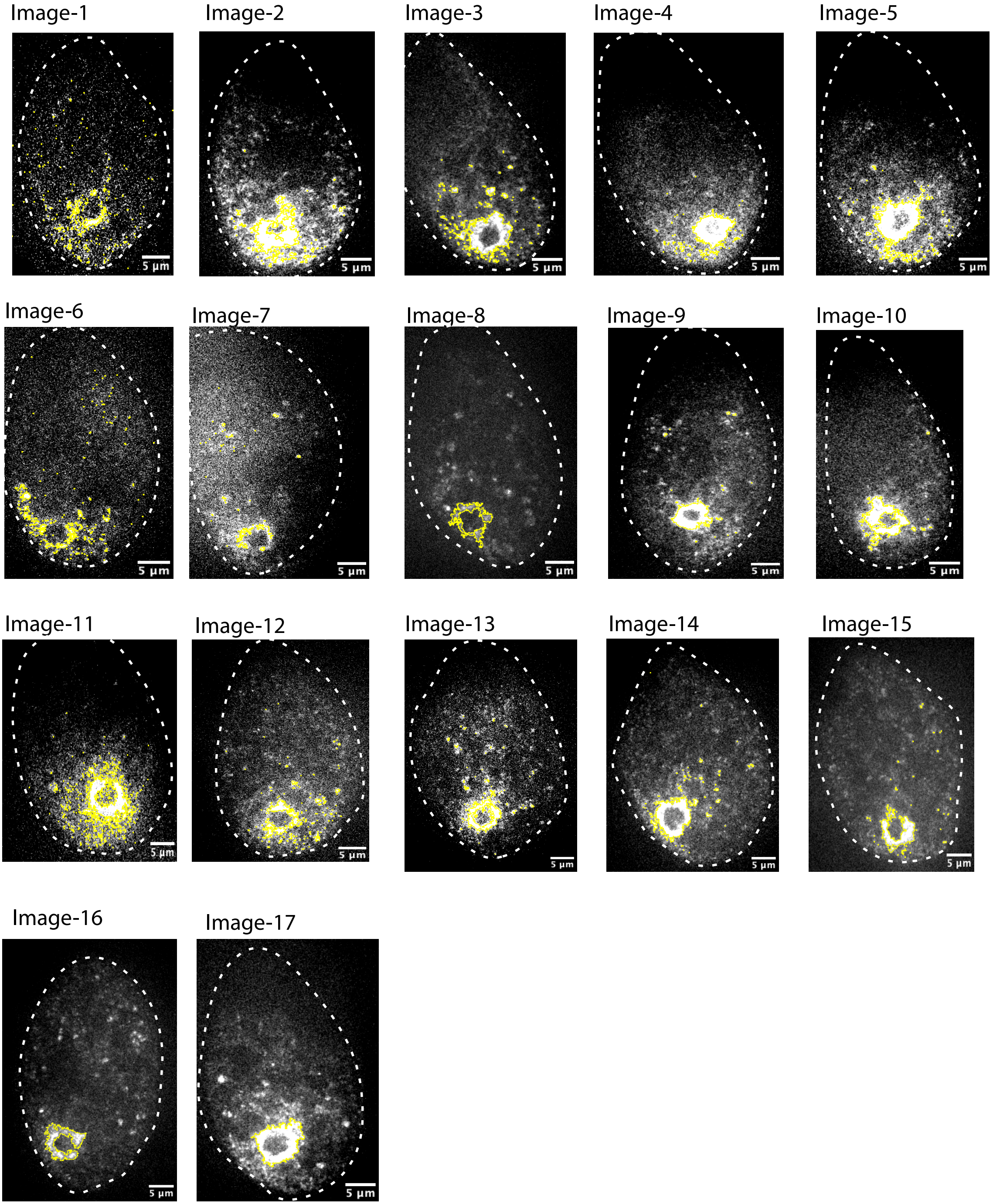

### Fig. S10

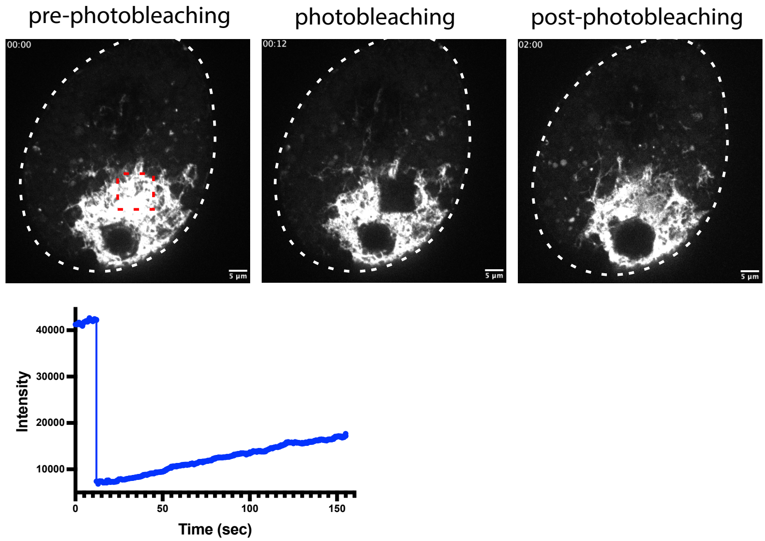

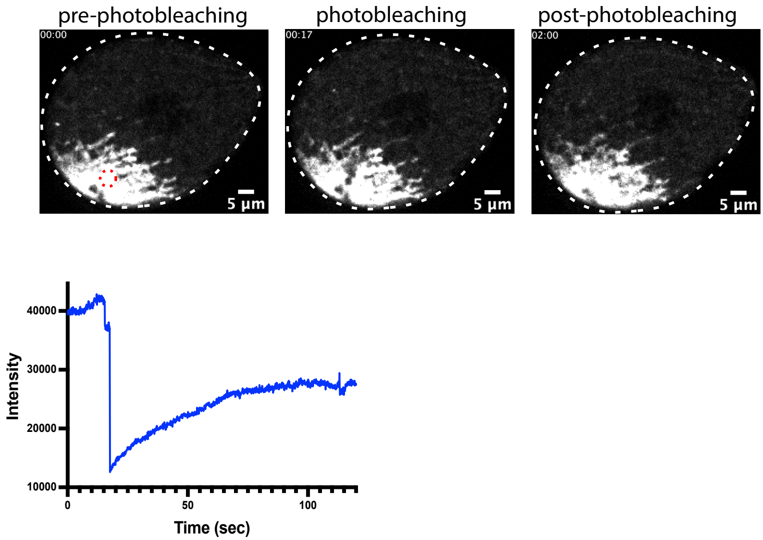

### Fig. S11

#### Fig. S11

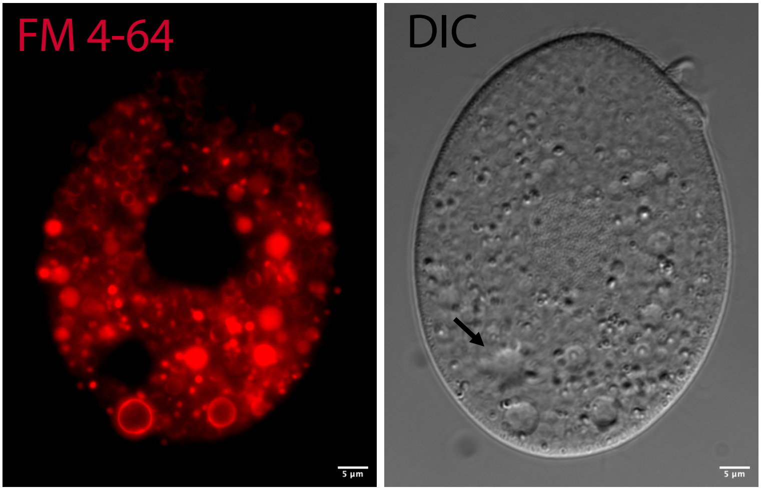
