## Supplementary material for "Structure and dynamics of the contractile vacuole complex in *Tetrahymena thermophila*": Fig. S2

**A**

| Protein ID | Enrichment | Intensity | Sequence coverage | Product Description |
|---|---|---|---|---|
| high confidence | | | | |
| TTHERM_00758980 | ∞ | 6.7E+11 | 82% | Dop1p |
| TTHERM_00637570 | ∞ | 8.1E+10 | 100% | Bis(5-adenosyl)-triphosphatase, HIT domain |
| TTHERM_00378790 | ∞ | 5.5E+10 | 38% | scavenger mRNA decapping enzyme carboxy-term-binding protein, N-terminal HIT domain |
| low confidence | | | | |
| TTHERM_00637780 | ∞ | 2.0E+09 | 2% | transmembrane protein, putative ion channel |

**B**

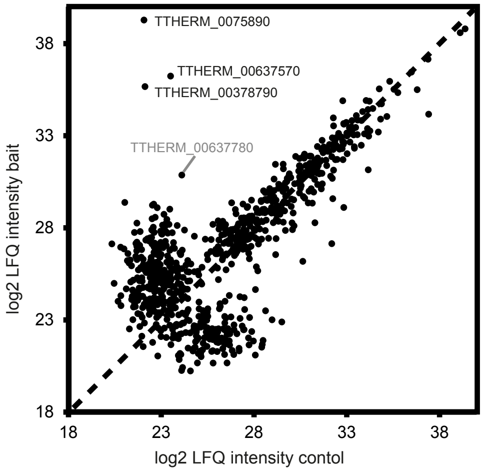
