## Supplementary material for "Structure and dynamics of the contractile vacuole complex in *Tetrahymena thermophila*": Supp Figure legends

Figure S1: Immunoprecipitation of Dop1p-interacting proteins.

A: A Coomassie blue-stained SDS-PAGE gel of the total cellular and anti-FLAG immunoprecipitated proteins from WT cells and Dop1p-FLAG-expressing cells. Lane 1 and 3: WT cells; lane 2 and 4: Dop1p-FLAG-expressing cells. Lanes 1 and 2: SDS whole cell extracts; Lane 3 and 4: pulldown eluates from anti-FLAG beads. IP=immunoprecipitates. Protein cryopowders were prepared from WT cells and Dop1p-FLAG-expressing cells and dissolved in 20 mM HEPES PH7.4, 250 mM NaCitrate. The pulldown eluate is enriched in bands near the 250 kDa marker (Dop1p~256 kDa with FLAG ~1kDa).

B: Western blotting of anti-FLAG pulldown. Samples identical to those in the SDS-PAGE gel from Panel A, with the equivalent lane loading, were used for Western blotting using the anti-FLAG antibody. The arrow indicates species in the expected weight range for Dop1p-FLAG.

Figure S2: Identification of candidate Dop1p-interacting proteins.

A: Table shows the top four most enriched proteins (including the Dop1p bait as the first one) identified by LC-MS/MS.

B: Multi scatter plot of the imputed logarithmic LFQ-intensity correlation comparing proteins detected in the FLAG-tagged Dop1p pulldown and the respective WT control. Protein cryopowders were prepared from WT and Dop1p-FLAG- expressing cells and suspended for incubation with anti-FLAG beads, as described in Materials and Methods. Bound proteins were eluted with 2X SDS sample buffer, electrophoresed a short distance into SDS-PAGE gels, and subjected to tryptic digests and LC-MS/MS analysis.

Figure S3: Live cell imaging of transmembrane protein TTHERM_00637780, identified in the Dop1p pulldown. TTHERM_00637780 was tagged with mNeon at the endogenous locus. The semi-regular array of cortical puncta resembles the pattern exhibited by known Golgi markers in these cells; in addition, other mobile puncta are visible deeper in the cytoplasm. Immobilized cells were imaged using a Zeiss Axio Observer 7 system. This image shows one frame from a time-lapse video (Movie 17).

Figure S4: Three additional images of the CVC in Vma4p-mNeon-expressing dividing cells. A and B: The newly forming CVC in the daughter cell becomes discernible as a Vma4p-labeled reticulum initiation, which seems to be interconnected with the parental CVC through a subset of cytoskeletal meridians. C: In the later stage of cell division (MAC amitosis), the developing Vma4p-labeled reticulum in the daughter cell becomes fully formed and exhibits a resemblance to the reticulum observed in the parental CVC. Growing cells were fixed as described in Materials and Methods. Images were taken with a Zeiss Axio Observer 7.

Figure S5: Live and fixed cell imaging of Scr7p (Scramblase 7).
A: SCR7 (TTHERM_00999060) was tagged with mNeon at the endogenous locus. Live images show that Scr7p localizes to the CV bladder (left panel) and the reticulum (right panel), in addition to dispersed puncta at the cortex and in the cytoplasm, particularly at the cell anterior, a localization shared by *T. thermophila* Scr1p (Chen et al., 2014). Images were taken with a Marianas spinning disc confocal microscope. These images show the selected frames from a time-lapse video (Movie 19)
B: Cells co-expressing Scr7p-mNeon and Vps8Dp-mCherry were fixed and imaged with a Marianas spinning disc confocal microscope. Shown are the separate channels and the merge, in which Scr7p and Vps8Dp tightly co-localize.

Figure S6: FRAP analysis of cells expressing Dop1p-mNeon. For each of three cells, the images show the cells before and after the photobleaching event, whose target in each cell is indicated with a dotted red line. The fluorescence intensities in the photobleached areas are shown in the right-hand chart. A: photobleaching focused on an area within the CV reticulum. B and C: photobleaching focused on the CV bladder. Live Imaging was using a Marianas spinning disc confocal microscope.

Figure S7: FRAP analysis of cells expressing Vps8Dp-mNeon. For each of three cells, the images show the cells before and after the photobleaching event, whose target in each cell is indicated with a dotted red line. The fluorescence intensities in the photobleached areas are shown in the right-hand chart. A: photobleaching of a part of the CV bladder. B: photobleaching of the whole CV bladder. C: photobleaching of the CVC in a focal plane where the reticulum is in focus. Live Imaging was using a Marianas spinning disc confocal microscope.

Figure S8: Images used to quantify Dop1p in the CV versus cytosol. The 17 images shown were analyzed by FIJI image process software to measure the signal sum intensity for the whole cell and for the CVC, indicated in yellow. Images were taken with a Marianas spinning disc confocal microscope.

Figure S9: Images used to quantify Vps8Dp in the CV versus cytosol. The 17 images shown were analyzed by FIJI image process software to measure the signal sum intensity for the whole cell and for the CVC, indicated in yellow. Images were taken with a Marianas spinning disc confocal microscope.

Figure S10: FRAP analysis of cells expressing Vma4p-mNeon. The images show the cells before and after the photobleaching event, whose target in each cell is indicated with a dotted red line. The fluorescence intensities in the photobleached areas are shown in the accompanying charts. Live imaging was using a Marianas spinning disc confocal microscope.

Figure S11: Intracellular labelling following long-term incubation with FM4-64. Cells were incubated overnight with FM4-64, and imaged with a Zeiss Axio Observer 7 system. Left panel: FM4-64 labels a wide range of intracellular structures but not the CV, whose position is indicated in the right-hand panel DIC image.  The CV could be identified unambiguously based on its characteristic contractile activity, as can be seen in the time-lapse video from which these images were taken (Movie 18A and 18B).

CHEN, B. C., LEGANT, W. R., WANG, K., SHAO, L., MILKIE, D. E., DAVIDSON, M. W., JANETOPOULOS, C., WU, X. S., HAMMER, J. A., 3RD, LIU, Z., ENGLISH, B. P., MIMORI-KIYOSUE, Y., ROMERO, D. P., RITTER, A. T., LIPPINCOTT-SCHWARTZ, J., FRITZ-LAYLIN, L., MULLINS, R. D., MITCHELL, D. M., BEMBENEK, J. N., REYMANN, A. C., BOHME, R., GRILL, S. W., WANG, J. T., SEYDOUX, G., TULU, U. S., KIEHART, D. P. & BETZIG, E. 2014. Lattice light-sheet microscopy: imaging molecules to embryos at high spatiotemporal resolution. *Science,* 346**,** 1257998.
