## Supplementary material for "Structure and dynamics of the contractile vacuole complex in *Tetrahymena thermophila*": Supp Materials and Methods

**Supplemental Materials and Methods**

**Endogenous expression of FLAG-tagged Dop1p, cryomilled cell powder preparation, FLAG immunoprecipitations and Mass Spectrometry.**

p-FLAG-ZZ-Neo4 was used to construct and generate a strain expressing endogenous Dop1p with a FLAG epitope at the C-terminus of the *DOP1* genomic locus. The initial Dop1p-FLAG-expressing transformants were serially passaged at least 20 times, while increasing the concentration of paromomycin from 120 μg/ml to 1000 μg/ml and decreasing the inducer concentration of $CdCl_2$ from 1 μg/ml to 0.05 μg/ml. Cryomilling was based on (Obado et al., 2016, Sparvoli et al., 2020). 10 L cultures of Dop1p-FLAG and CU428 were grown to 2–5 x $10^5$ cells/ml, washed once with 10mM Tris-HCl pH 7.4 and re-pelleted. Supernatants were rapidly aspirated to leave a dense cell slurry. The slurries were dripped from a volumetric pipette into liquid nitrogen, and the frozen beads collected and milled to powders using a Cryogenic Grinder 6875 Freezer Mill, and stored at -75°C.

Pulldowns were as described previously (Sparvoli et al., 2020): 10 gram of cryomilled cell powder was weighted and immediately suspended in 50 mL dissolving buffer containing 20 mM HEPES PH7.4, 250 mM NaCitrate on ice, supplemented with protease inhibitor cocktail tablets (Roche), gently mixed by inversion for 1h at 4°C, and then on ice until no solid matter was visible. The solutes were then centrifuged at 140k x g (Beckman Instruments type 45 Ti rotor) for 1.5h at 4°C, and the supernatants transferred to new tubes, for incubation at 4° C for 2 h with anti-FLAG beads (EZ view Red Anti-FLAG M2 affinity Gel, Sigma) which had been pre-washed with lysis buffer for 2 hours at 4°C. The beads were then washed five times with 20 mM Tris-HCl pH 7.4, 1 mM EDTA, 500 mM NaCl, 0.1% NP-40, 1 mM DTT, 10% glycerol supplemented with protease inhibitor cocktail tablets (Roche) and eluted with 60 μl of 4X LDS sample buffer (988 mM Tris, 2.04 mM EDTA, 8 % LDS (Lithium dodecyl sulfate), 40 % Glycerol, 0.88 % Commassie Brilliant Blue G250, 0.7 mM Phenol red) + 40 mM DTT for further SDS-PAGE electrophoresis. Protein samples were electrophoresed on a 4–20% gel SDS-PAGE gel to verify pulldown efficiency by Coomassie blue staining and Western blotting using anti-FLAG antibody. The eluted sample (total in 60 µl sample buffer) was used to run a SDS-PAGE, with the dye front allowed to electrophorese ∼1 cm into the gel. The gel was then stained with Coomassie Blue R-250 solution (0.1% w/v Coomassie, 10% acetic acid and 50% methanol) to visualize the lane. A single 1 cm gel slice was excised and prepared for protein identification via mass spectrometry.

**Mass Spectrometry**

Immunoprecipitation eluates were loaded on a 4–20% gel for SDS-PAGE gel, allowed to migrate for ∼1 cm into the gel, and briefly stained with Coomassie Blue R-250 solution (0.1% w/v Coomassie, 10% acetic acid and 50% methanol). The protein containing gel slice was excised from the Coomassie-stained gel, destained, and then subjected to tryptic digest and reductive alkylation using standard procedures. LC-MS/MS was performed by the OMICS Proteomics Facility at Biocev on a Ultimate3000 nano rapid separation LC system (Dionex) coupled to a Orbitrap Fusion mass spectrometer (Thermo Fisher Scientific). Mass spectra were processed using the intensity-based label-free quantification (LFQ) method of MaxQuant version 1.6.6.0 (Cox and Mann, 2008, Cox et al., 2014) searching the *T. thermophila* annotated protein database from ciliate.org (Eisen et al., 2006, Stover et al., 2006). The minimum peptide length was set at seven amino acids and false discovery rates (FDR) of 0.01 were calculated at the levels of peptides, proteins and modification sites based on the number of hits against the reversed sequence database. If the identified peptide sequence set of one protein contained the peptide set of another protein, these two proteins were assigned to the same protein group. Potential interactor were ranked by enrichment ratios comparing to a control sample and respective LFQ intensities. Proteomics data have been deposited to the ProteomeXchange Consortium via the PRIDE partner repository (Perez-Riverol et al., 2019) with the dataset identifier PXD043302.

**Western blotting**

Total cell protein extract and immunoprecipitated samples were analyzed by Western blotting as previously described (Sparvoli et al., 2018). Proteins were resolved with the

Novex NuPAGE Gel system (8% or 4–20% Tris-Glycine gels, Invitrogen), and transferred to 0.2 µm PVDF membranes (Thermo Scientific). Blots were incubated with anti-FLAG (1:2000 dilution, Sigma) antibodies for overnight at 4 °C after pre-blocking with 3% bovine serum albumin in TBST (1X Tris-Buffered Saline, 0.1% Tween® 20). Secondary antibody conjugated with ECL Horseradish Peroxidase (NA931) (GE Healthcare Life Sciences, Little Chalfont, UK) was diluted 1:20000 for incubation with the blots, and signal was visualized by SuperSignal West Femto Maximum Sensitivity Substrate (Thermo Scientific).
