## Supplementary material for "Structure and dynamics of the contractile vacuole complex in *Tetrahymena thermophila*": Movie Legends

Movie 1: Live cell expressing Dop1p-mNeon with cross-sectional view of the CV. Linked with Figure 1C. Live imaging with a Marianas spinning disc confocal microscope.

Movie 2: Live cell expressing Dop1p-mNeon, with focal plane near the top surface of the cell. Linked with Figure 1D. Live imaging with a Marianas spinning disc confocal microscope.

Movie 3: Live cell expressing Dop1p-mNeon, at focal plane very near the plasma membrane. Linked with Figure 1E. Live imaging with a Marianas spinning disc confocal microscope.

Movie 4: Live cell expressing Dop1p-mNeon, at focal plane corresponding to the CV cross-section. Linked with Figure 2A. Live imaging with a Marianas spinning disc confocal microscope.

Movie 5: Live cell expressing Dop1p-mNeon, at focal plane tangential to the CV. Linked with Figure 2B. Live imaging with a Zeiss Axio Observer 7 system.

Movie 6A: Live cell expressing Vps8Dp-mNeon, in the DIC channel. Linked with Figure 4A. Live imaging with a Zeiss Axio Observer 7 system.

Movie 6B: Live cell expressing Vps8Dp-mNeon, in the fluorescence channel. Linked with Figure 4B. Live imaging with a Zeiss Axio Observer 7 system.

Movie 7: Live cell expressing Vps8Dp-mNeon, showing Z-sections from the cell surface to the interior. Live imaging with a Zeiss Axio Observer 7 system.

Movie 8: Live cell expressing Vps8Dp-mNeon, at focal plane corresponding to the CV cross-section. Linked with Figure 4C. Live imaging with a Marianas spinning disc confocal microscope.

Movie 9: Live cell expressing Vps8Dp-mNeon, at focal plane tangential to the CV. Linked with Figure 4D. Live imaging with a Marianas spinning disc confocal microscope.

Movie 10: Live cell expressing Vma4p-mNeon in the side view of the CVC near the plasma membrane and proceeding with different Z-stacks between cell interior and cell surface. Linked with Figure 5C. Live imaging with a Zeiss Axio Observer 7 system.

Movie 11: Live cell expressing Vma4p-mNeon, at focal plane tangential to the CV. Linked with Figure 5D. Live imaging with a Marianas spinning disc confocal microscope.

Movie 12: FRAP analysis of cell expressing Dop1p-mNeon. Linked with Figure 8A. Live imaging with a Marianas spinning disc confocal microscope.

Movie 13: FRAP analysis of cell expressing Vps8Dp-mNeon. Linked with Figure 8D. Live imaging with a Marianas spinning disc confocal microscope.

Movie 14: FLIP analysis of cell expressing Vps8Dp-mNeon. Linked with Figure 8G. Live imaging with a Marianas spinning disc confocal microscope.

Movie 15: FRAP analysis of cell expressing Vma4p-mNeon. Linked with Figures 9A and 9C. Live imaging with a Marianas spinning disc confocal microscope.

Movie 16: FRAP analysis of cell expressing Dop1p-mNeon, focusing on CVC reticulum. Linked with Figure 9D. Live imaging with a Marianas spinning disc confocal microscope.

Movie 17: Live cell expressing TTHERM_00637780-mNeon. Linked with Figure S3. Live imaging with a Zeiss Axio Observer 7 system.

Movie 18A: Cell incubated overnight with FM4-64, viewed in the red fluorescence channel.  Live imaging with a Zeiss Axio Observer 7 system.

Movie 18B: Cell from Movie 18A, viewed in the DIC channel. Live imaging with a Zeiss Axio Observer 7 system.

Movie 19: Live cell expressing Scr7p-mNeon. Linked with Figure S4. Live imaging with a Marianas spinning disc confocal microscope.

Movie 20: Cells expressing Scr7p-mNeon. Live imaging with a Zeiss Axio Observer 7 system.
