## Supplementary material for "Structure and dynamics of the contractile vacuole complex in *Tetrahymena thermophila*": Supp Table caption

Supplemental Table caption:

Table S1: Cell strains used in this study.

Table S2: PCR primers used in this study.

Table S3: Quantification data for Dop1p and Vps8Dp.

Table S4: Mass spectrometry data-1.

Table S5: Mass spectrometry data-2.
