## Supplementary material for "Structure and dynamics of the contractile vacuole complex in *Tetrahymena thermophila*": Table S4

| **Protein ID** | **Enrichment** | **Intensity** | **Sequence coverage** | **Product Description** | **TMHMM** | **SignalP** |
| --- | --- | --- | --- | --- | --- | --- |
| high confidence | | | | | |  |
| TTHERM_00758980 | ∞ | 6.7E+11 | 82% | Dopey | 0 | 0 |
| TTHERM_00637570 | ∞ | 8.1E+10 | 100% | Bis(5-adenosyl)-triphosphatase, HIT domain | 0 | 0 |
| TTHERM_00378790 | ∞ | 5.5E+10 | 38% | scavenger mRNA decapping enzyme carboxy-term-binding protein, N-terminal HIT domain | 0 |  |
| low confidence | | | | | |  |
| TTHERM_00637780 | ∞ | 1.95E+09 | 2% | transmembrane protein, putative ion channel | 10 | 0 |
| TTHERM_00194540 | ∞ | 6.98E+08 | 17% | TATA-binding protein interacting (TIP20) protein | 0 |  |

>TTHERM_00637570	Bis(5'-adenosyl)-triphosphatase

MINTALKLGSIEIPQQMIFWTKSNICAIIPCVQLVPGHVLIIPKRNVSYFNDLELQEVFD

IGLLTRFLTKGLEKFYTATSSTVYIHNYNPNDSESLQQVYVHIIPRKPADFQNNDDIYKK

LEEYDAEFTKKFKWGFTQANSSLNGVLEIEANECKKYATFLETFQREEAEKS

>TTHERM_00378790	scavenger mRNA decapping enzyme carboxy-term-binding protein

MSASVVANLTVQFGKSQLPLSQIFILRKNVFATTNLKPACPGHVLVASRRPVKRLHELTE

VETLDLWTTVQQVSRVMEQIHKFPCQIGVQDGTDAGQTIDHVHIHIIPFPKEYSQDVIMD

SEGRIKCNLNEENQIQKYYQIQKDLLEQQMIWPKKLTNIELTLNKGQTSSIDQIQIDQSI

QLNSFINVNNIFLDDICSYVFRQCNQFILKYYALQKYDSANQYIDKQIDILCLKYMISKI

SIVDKHLHIIFYIKISLKLQQEGQLKVDRLFVCLFD

>TTHERM_00758980	dopey, amine-terminal domain protein

MDKTKQIEKSSQLNKQLNTLLSGFEKNKAWSDIGTWLYKVEQAMKENPSPYISEKLSLAK

RLAQCLNPVLPQAINLQTLKIYELIFENIKIASQGNVEEYKKMFCEDIGIYSVGLFPFFQ

NAHQKNKHQFLEIIRNYYIPVGRDLIPCLPGLVLSILPGLEEQSEILIKDVTKTLEAIMT

SCGRRYFLGSVWMGIKRTQRGREHAIKFLTKLLPKVKYEEDEEENQDASDNSSNFSEDNY

EEFDETKFKEQQKKEQTQTVEQTPVKQNQSPKFQEQQAEQADQVDHDQQNNQVEIQDEEK

IAQDLIQQAQAEAEKHFQSMLEGNEENQDQQEQHQEEQQPQQPVEEQEVEDDQQQKQLNR

QQADEEIQKSLVASLQQQQQLQQQQQQLQQQQQQQQQQQQQQQQQQAATSTSTAPVSAAK

SLTDQKLNILESTSEQKKGGKSALEIAFQHEIAPKIAKKQDFTNTFFVRDNLKYIEESIN

ENDDNNNFPNKSSLIINTILSCFEDENQIVRRNILDLMYHHLRINHRILSRDDQLILVEG

VLYLLIRKDVSVTRRVNVWLFGKPDMDNKYTITKSREFVVDLLIDAFKRIFSVEPQDKVA

ALQPIKIMQQFYMEHENLVEITLSKLAISFVKYVYTHTQNMQFSADVLKSGIRFMENIQG

QLNSLLKSISEEAVQLLREKKDDQCLEIINLIEFTYENLLASEDSFDPMQQRDCIKETLS

RLFISLSYLDEKNIQEFHSLEPTLKVIYKLMLKLESAEAKIKAIEKKEKKQLEITNEQKE

IAQYLDECSEKYFNFYGKLSNELENFKSNNIYKLASQSLVKLQKFLFDKKKMTQLPQWFK

CISQSIKITNPNLSLIAIESMIEIMISEKIDPIYEQLKVLIIDEAKLKYSKKNELVAGND

YTKLTLEKLWSLLDFQHFYDRIIDLIIYFSKYFPQYFKEVVVNSFQAVSITEKEASIRRF

AVFWRLTAHHKSKLLLSELNKVGLFIMLDFLDHDNPLIRHASKNWLMESVPQFYRIIDPL

FEVLLQANSSWYVTDSYQVFYTKIYETTRANETFRKLKSILIIASDVFLKYISSYQLSPL

LQDMKSHFTDKNQAVLLRKLGATMSYFSTINPNLNSLDMTSSQSTAVNDKKVTDLKMTYL

DLLIIVCLRYIQGQALESLSLKFQVENAAVNASACEYMELLITHLEDAPLCLALCEYLME

PLLTILGHCITNRDYVMQVQLLNVFRIIFFHSSYIKKNNTEEIKFKLVQTLSNKLFIPNI

LKGLMTSIPYVRSQFISFISACIPLLSTYLASNQLTSIIKSIMFTYFSLIKHLATTPAQD

NDESMVTISKTQIIKQPSLIKQNKTVILIQTQEVVKEDSNIYEIHTVLEGVRTILQFFLK

IRTLEEIESSKKDQSGGFANLMKTVLTLGLAGSGNANDKDKDGKAFFEKYPDTCKSLLYD

IKQVVELFINSWQTTPEFITEFTSYGIRPYEQKKFNSFNNLLMDMIKNAENDKQRVKILI

ISIIKPLVYEFPNDMMNSFLLLWNRSCMTDPPNAHKDFNQNEFLKKMIEMITILNIPNEL

FLESFYQSFISEIIQKFYQQKNTKEKKNVYFLNYEIAQHESKLLYFLYIYLSYTYIDSNI

LKKENLLLFWNIILKILKIFQPSKNPNTVMWMLDLLYMLSEKYSPKEILSDSKFKKELHD

LINEKLSYLSGWIANVYQVKFNEPSAQTQQSLNNAVDQHMKEMIVQSLQQYRIISPYSPT

FYEISLSFYSQIHILEHAPKSIANQMKELQMNQLDQFIFDRIDTVDENAVYDKYRLICLE

TLKTLSLELLQNTYSPERQDRVVVRVKEFVDPLYAVIENKGSTNSYFLESTSELICSLIE

KAPSLLIKDFRKSILEMFNKDNFFSCNKGTLRYWGKIIDYVLTQDKSNDLFNEYLEKVSL

SAGFFAREQNENKKKIKAFKRTCFIIFSGAKDKYANRLRKLLDSIAEVIKTTDSSQSSLL

ILILFCIRILILRLSQSNLNELFRQIWPILLTLLIQIYSKPLPGQPPRNPNLMLAGLKLI

EILSVAQLDEFFLHQWMFIFDYFGLKLQPLSQQERENQIRLRGQMNEAQLRQYAKISPFY

FQPFVSTCLNEQDLGVKYVNQPIDESEQSEQITYQKKQRKLIMTQCKVDDENELKSHALS

LCQYLISNNENRTEIDFNSIEALIEDDFINLDEHILKV

>TTHERM_00637780	transmembrane protein, putative

MSQQKVIAQQMSLRKQVMAKSKTLFQAEKAILDQEKVSDQLASWEEAVAYRNKFSVMKNN

GGSYEEYSLNCDLSELAQYGPGLRLFFELLKYSAIVFLLMSLISIPALIGNIQGQSLNSI

EMSTNSLPKLSISNQPKLKLLSPGANASDADYQADQVENQQRIVDFVSNSDSRLIQVAVP

DIVYSVIFLLFIFWFHYHSHQIAKETDLKNSLPSNYSIEVSGFPHTITDEKILSNHFRDN

FSVDVFECKFARNYYNTLFLHKEEAQLIDLIKQEKSRQQFLGKDEKKSETLQKLGKKLAD

IQADINQQVQTKMNGSVLSHNEYPSVKAFIILNSIEDKQKIQQEYKKTKGFFGQKRVKEF

MLQNKHLLKLNFKPDEPSNILWENLEVSSFNRFLRTLAVIILVIIVMIITFAVIIIANIA

TPQNTEDCPKQSISFSQASKSSLYTQCYCTNVSFSQMTSNSALQNLCWDVWVKYATTYAL

TIVTGLVVLIVNFLLRLTIQALGKFNRYKTITKYTTSMTSKLFLAFFVNTALITLFLQAN

IYGFVPAITFSKPIPPISNLQDNNKSQFSTDFDRSWYLQVGSKITVTVFFFVVTLFLFQF

LYSMISKCRRESKVKQLEGKDVQRVANKTILGPEFPISFYYASTFNIIFTTLFYSSGIPI

MLFAGFIILTSQYWVYKYLLLRVCRRPPTYDTGLSNRMLLILPWSLLLHLAVGLYMYGQP

LIFPSSQSQLVLNMNQTTGEVKVKINDSAPIENRAFHILYLFIFLLIVLGLYILNISYSG

FLQRFVNMCCKRSHQVVPQHQLVPYNEEYFNIEKRMLPSYNIKVNQDYRYIIEAIDLEKK

FDSENSPRANSSSPQYKETDSINKNIDSIIENI

>TTHERM_00194540	TATA-binding protein interacting (TIP20) protein

MQSAQQQQQQLLDQKRFDELVQDAKDFDPDKKFMAANDLSNALSNGQLKDTDYDTVVKIL

LNQLNDDSNEVQGNSIRCLSKIINKLQETQVENVSKTMIVKVIEDKGEYRDVYATCLKTL

INGVPSNYAKAVQPILLHAIQGMEQKLNNKEFDVQEELCDILNNLFKKWGQSLSTTNANV

KNLTNILMSNISKGERNSLKKKSCSCLGSLGLILHKDDIGRVVKELLQNIKKPDNQKQIQ

QHLREKQIYVVYALSAISKTVAKKLGSNLKDLINIILKDINQYVEEVDDYDLIITISEMF

ESYLSILESLIKGCPDEIKEYFPQIVDLSSQLINYDPNGQARISNGGEMEIEDGGEEDEY

DYLSDDQDGDDSSWRVRRAALNIIETLVKSDSEMIRPIFEKCVSSQSEQSTIVYRLGERN

ENIRFATFSCLQTIIRAIVISDVSHDKGDDDFDQELNSGPQLVRVKSAVKEFNISEIISE

IFNTLTFILEKEKKVTQAILTGAYKLIESISRNLSRQVVSDPSIWRLAFDLIKRTIENKA

SPNELKVTAFSSLRKLLKNYHESSEIAHLQNDIPNIFKLASVGLVQDYFKIVAESFRCLN

SLFNMINKQFNSAIPTFSASIQQIFPQISVKLNSSDIDQEIKQSVLSCTTSLLVSLPQTI

QAQQIKDIINAISEKIKSETKKIPVLKLLKKIPLSVHPHITADSIKLLNSSAESICDLLA

KQDRSLRVASLEALDKTLVLLSEKKSPLSAKARDNFIALAYDFLNDKDTWVAQLYLSVLK

KSIENAPQSPQYYTKIIESITKFSNSSIVSSAHFSSLYEVFATLSKLNLSNRSQMIQNLI

ATGSKDSINSVSRAISSIIIELPAGEQQTFINQIFSTASSDKEVVLKRQICLVTLGEIGR

HVNLCSGNILQEVQKILNNKQNNEEIRTAASISLGGIAIGNLNMVLPQVIQTINSGSEGQ

YLMLNSLKQIIEHSQQLSQSIIQIIPNLFKATENGDESLCNIISYIIGKVTHLNLKEMKP

LILQNLSSKNQNTKYTTASSFKYFCLKTLKIDNDLRELVYALMNNINEKDIRTRTAILKS

LNMISYNLPNAIIQHINKNEFFLPVREALRFTQIREIDFGPFKQKNDDGEPVRNAAFTLL

DTAIDHLHYRGVENFEREILEEVLYEFANESSEDIKILRFQICTKFAHKVPLKVTPFLDQ

ISEVFLNVLKPYASKADGDRAADMVRTGLICCLTLKSIAEDESNTKYLNFWELVMKTEKY

RNIINSMTN
